# Cellular response to small molecules that selectively stall protein synthesis by the ribosome

**DOI:** 10.1101/461624

**Authors:** Nadège Liaud, Max A. Horlbeck, Luke A. Gilbert, Ketrin Gjoni, Jonathan S. Weissman, Jamie H. D. Cate

## Abstract

Identifying small molecules that inhibit protein synthesis by selectively stalling the ribosome constitutes a new strategy for therapeutic development. Compounds that inhibit the translation of PCSK9, a major regulator of low-density lipoprotein cholesterol, have been identified that reduce LDL cholesterol in preclinical models and that affect the translation of only a few off-target proteins. Although some of these compounds hold potential for future therapeutic development, it is not known how they impact the physiology of cells or ribosome quality control pathways. Here we used a genome-wide CRISPRi screen to identify proteins and pathways that modulate cell growth in the presence of high doses of a selective PCSK9 translational inhibitor, PF-06378503 (PF8503). The two most potent genetic modifiers of cell fitness in the presence of PF8503, the ubiquitin binding protein ASCC2 and helicase ASCC3, bind to the ribosome and protect cells from toxic effects of high concentrations of the compound. Surprisingly, translation quality control proteins Pelota (PELO) and HBS1L sensitize cells to PF8503 treatment. In genetic interaction experiments, ASCC3 acts together with ASCC2, and functions downstream of HBS1L. Taken together, these results identify new connections between ribosome quality control pathways, and provide new insights into the selectivity of compounds that stall human translation that will aid the development of next-generation selective translation stalling compounds to treat disease.

## INTRODUCTION

Proteins are the main druggable therapeutic targets for the treatment of human diseases, ranging from metabolic disease, to cancer and dementia [1]. Most therapeutic strategies consist of inactivating key protein enzymatic or binding activities by using small molecules or antibodies. However, many proteins remain “undruggable” due to the difficulty in identifying small molecule or biological inhibitors of their function [2]. An alternative approach would be to prevent the synthesis of target proteins in the first place, by targeting the step of translation [3]. However, targeting translation in a highly specific way remains an unsolved problem. Many known translational inhibitors bind to the ribosome in close proximity to messenger RNA (mRNA) or transfer RNA (tRNA) binding sites, and thereby interfere with different steps of the translation process [4]. These mechanisms of action, even if context-specific, lead to substantial inhibition of translation which is extremely toxic, as exemplified by the many classes of antibiotics that target the bacterial ribosome [5,6]. In humans, the low specificity and high toxicity of homoharringtonine (HHT), the only translational inhibitor approved as a drug treatment, makes this compound useful as a last resort treatment for chronic myeloid leukemia (CML) [7,8], but it is not clear how its mechanism could be repurposed for specific drug targets.

As opposed to affecting translation for much of the proteome, a new class of compounds has been identified that selectively targets the translation of PCSK9, with few off-target effects [9–11]. These compounds bind in the ribosome exit tunnel [12], and seem to affect the trajectory of the growing nascent polypeptide chain in the exit tunnel as it is extended, thereby selectively stalling translation of a narrow spectrum of transcripts. Understanding the molecular basis for how these compounds selectively stall translation will be critical for the future design of transcript-specific translation inhibitors to target previously undruggable proteins [2]. However, designing new molecules as selective translation inhibitors to treat disease also necessitates a better understanding of how cells respond and adapt to compound-induced translational stalling.

Genomic screens serve as powerful and unbiased tools to identify genetic modifiers of the action of small molecule inhibitors. They can be used to discover proteins involved in an inhibitor’s mechanism of action, and to assess possible targets for combination therapy [13,14]. In particular, CRISPR interference screens (CRISPRi) allow robust and highly specific knockdown of gene transcription with minimal off-target effects [15]. These screens can be used to compare the effects of inhibitors on cell growth as a function of genetic background, and to identify components of genetically related pathways and how these are influenced by drug treatment [16]. Here, we used a genomic CRISPRi screen using a compound that selectively stalls the translation of human PCSK9, to test the effects of this class of compound on human cell fitness. We used compound PF-06378503 (PF8503) [9], a compound related to the PCSK9 selective translational inhibitor PF-06446846 (PF846) described previously [10], but which is slightly more toxic, to exert the selective pressure required for growth-based CRISPRi screens [17]. We show using ribosome profiling that PF8503 inhibits translation of an overlapping set of proteins compared to PF846, along with a distinct set of off-target proteins. In the CRISPRi screen, we identified proteins that suppress or enhance the toxicity of PF8503, a number of which are associated with translation and ribosome quality control pathways. We used targeted CRISPRi to validate the involvement of these proteins in cell fitness during PF8503 treatment, and to identify genetic interactions between ribosome quality control pathways. We also compared the genes identified in the PF8503 CRISPRi screen with those identified using the non-selective translation inhibitor HHT, in order to reveal the pathways likely to be general cellular responses to the stress induced by translation inhibition. Taken together, these results reveal pathways affected by selective stalling of translation, and suggest the cell dependencies most likely to be impacted by this class of inhibitors.

## RESULTS

### Cell toxicity of PF8503 and PF846

Growth based CRISPRi genetic screens require variation of cellular fitness of knockdowns to identify pathways and proteins involved in resistance or sensitivity to a stress condition. Therefore, to carry out a growth-based screen, a slightly toxic concentration of compound must be used [17]. Previous studies showed that cells from the hematopoietic lineage are more sensitive to the class of compounds that include PF846 and PF8503 (Fig 1A) [9]. In these experiments, which used rat bone marrow, PF846 and PF8503 had similar toxicity profiles [9]. The CRISPRi screen library was originally developed and validated in K562 cells [15], immortalized leukemia cells that can be induced to develop characteristics similar to early-stage erythrocytes, granulocytes and monocytes [18]. Given their close relationship to the hematopoietic lineage, we therefore used K562 cells to implement the CRISPRi screen. We first compared K562 cellular viability as a function of PF8503 and PF846 concentrations, and found PF8503 had a slightly higher negative impact on cellular metabolic activity and was slightly more toxic than PF846 (S1 Fig). We determined the best concentration of PF8503 to use in the CRISPRi screen to be 7.5 μM, which decreases cell viability 30-40% (S1 Fig).

**Fig 1.**
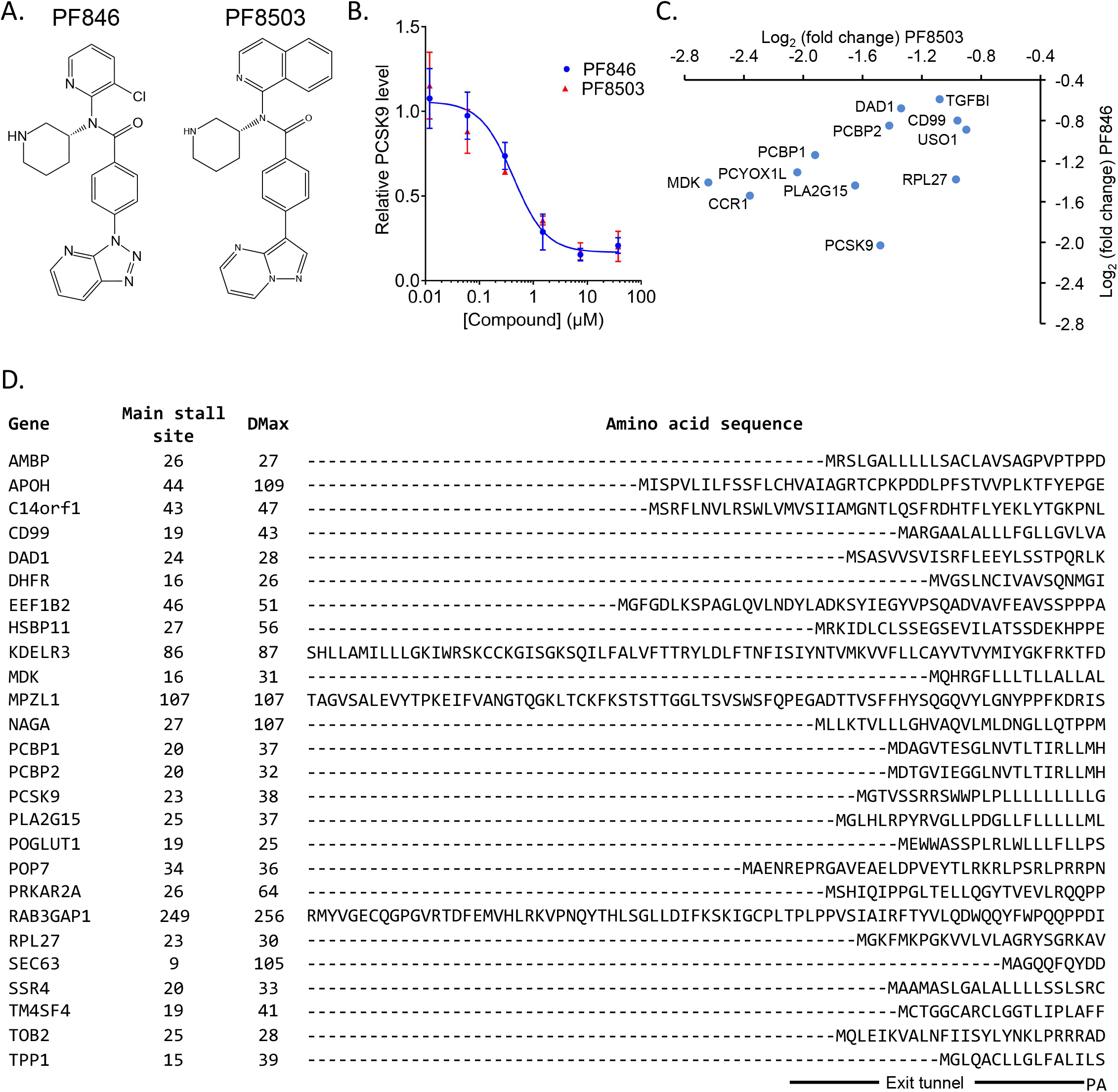
Comparisons of translation inhibitors PF8503 and PF846. (A) Structures of PF-06446846 (PF846, left) and PF-06378503 (PF8503, right). (B) Effect of PF846 and PF8503 doses on extracellular levels of PCSK9 secreted by Huh7 cells after overnight incubation. Experiment carried out in biological duplicate, with standard deviations shown for each compound concentration. (C) Proteins stalled by PF8503 or PF846 in Huh7 cells after 1 hr treatment, shown based on the log_2_(fold change) of reads relative to vehicle control. Reads were quantified 3’ of the DMax position, as described in the Materials and Methods. Experiment carried out in biological triplicate, with the average plotted. (D) PF8503 sensitive protein sequences in Huh7 cells. The sequences are aligned according to the last main pause site position. Putative locations of the nascent peptide chains in the exit tunnel and relative to the ribosomal P and A sites are marked. Also indicated are the codon positions of the stall site and DMax positions.

### Ribosome profiling to identify proteins targeted by PF8503

Before using PF8503 in a CRISPRi screen, we first used ribosome profiling to compare proteins targeted by PF8503 and the previously described compound PF846 in liver-derived Huh-7 cells, which endogenously produce PCSK9 [10]. We first treated Huh-7 cells with increasing concentrations of PF8503 to identify its IC50 value with respect to inhibiting PCSK9 production, which was similar to that of PF846, or ~0.4 μM (Fig 1B). We then treated Huh-7 cells with 1.5 μM PF8503, which corresponds to ~70% of the maximal inhibition of PCSK9 production, or 0.5% DMSO (vehicle control) for 1 hour prior to isolating ribosome protected fragments for ribosome profiling library generation. The same experiment and pipeline was used in parallel with PF846 in order to compare our results to previously published data [10]. After using the bioinformatic pipeline described previously [9] (S2A Fig), we found that PF8503 affected the translation of 46 mRNAs, whereas PF846 affected translation of 24 mRNAs after 1 hr of treatment. As expected PCSK9 was affected by both compounds with a log_2_ fold change of −1.5 and −2 for PF8503 and PF846, respectively (Fig 1C and **S1 Table**).

The percentage of mRNAs affected by PF8503 (0.50%) is comparable to that for PF846 (0.27%), (S3A Fig). The protein targets of PF8503 overlap with those of PF846 (10 of 24 proteins, Fig 1C), but PF8503 and PF846 inhibit the translation of 36 and 14 distinct mRNAs, respectively (**S3B** and S3C Fig). Notably the potency of PF8503 stalling on many of the common targets is higher than that of PF846 (Fig 1C and S3 Fig), as assessed by differential readcounts 3’ of the stall sites (defined by DMax) (S2 Fig). Furthermore, some of the additional proteins stalled in PF8503-treated cells are slightly impacted in the PF846-treated cells, but to a lower extent that did not allow these to pass the statistical filters (S3C Fig and **S1 Table**). We found that the stall sites on a given mRNA occurred at identical or nearly the same codons of the transcript when comparing ribosomal footprints from the PF8503- and PF846-treated cells. As observed previously, stalling occurs near the N-terminus of the protein, although not exclusively (Fig 1D and S4 Fig).

We also compared our present PF846 results with the 21 PF846 targets previously identified in Huh-7 cells (S3D Fig and **S2 Table**) and observed that changes in the read count 3’ of the stall site defined by DMax were highly correlated between experiments (Pearson R=0.82), showing the high reproducibility of our method of analysis (S2 Fig). However, we found that the main stall site position seen in the ribosome profiling read-counts varied slightly when comparing the present PF846 treatment and the previously published PF846 data [10], and when comparing PF8503 and PF846 stall sites on common targets (S4 Fig, **S1 Table** and **S2 Table**). This could be due to technical variability in the preparation of the ribosome profiling libraries, such as the efficiency of RNAse I digestion. Alternatively or in addition, this variability may reflect that compound-induced inhibition of translation is not due to a sharp stall at one codon position, but occurs due to a slow-down of translation over multiple adjacent codons [12]. Altogether, these results indicate that PF8503 is a selective inhibitor of translation and shares a similar mechanism of action to that of PF846 in stalling protein synthesis.

### CRISPRi screen to uncover genetic modifiers of cellular growth in the presence of PF8503

To identify pathways involved in the cellular response to high concentrations of PF8503, we used a genome-wide CRISPRi screen with an established whole-genome library of sgRNAs [17]. Human K562 cells constitutively expressing dCas9-KRAB-BFP were cultured with 7.5 μM PF8503 or 0.5% DMSO as a control. We used deep sequencing of the genomically-expressed sgRNAs in the cell population at the start and end of the experiment to determine their enrichment or depletion [17]. Briefly, sgRNA enrichments in the DMSO-treated control and PF8503-treated populations correspond to growth phenotypes termed Gamma and Tau, respectively, and quantify the impact of sgRNA expression on cell fitness independent of treatment. The difference in sgRNA enrichment between the control and PF8503-treated population corresponds to the impact of the compound on cell fitness independent of the impact of sgRNAs on cell growth, a parameter termed Rho (Fig 2A).

**Fig 2.**
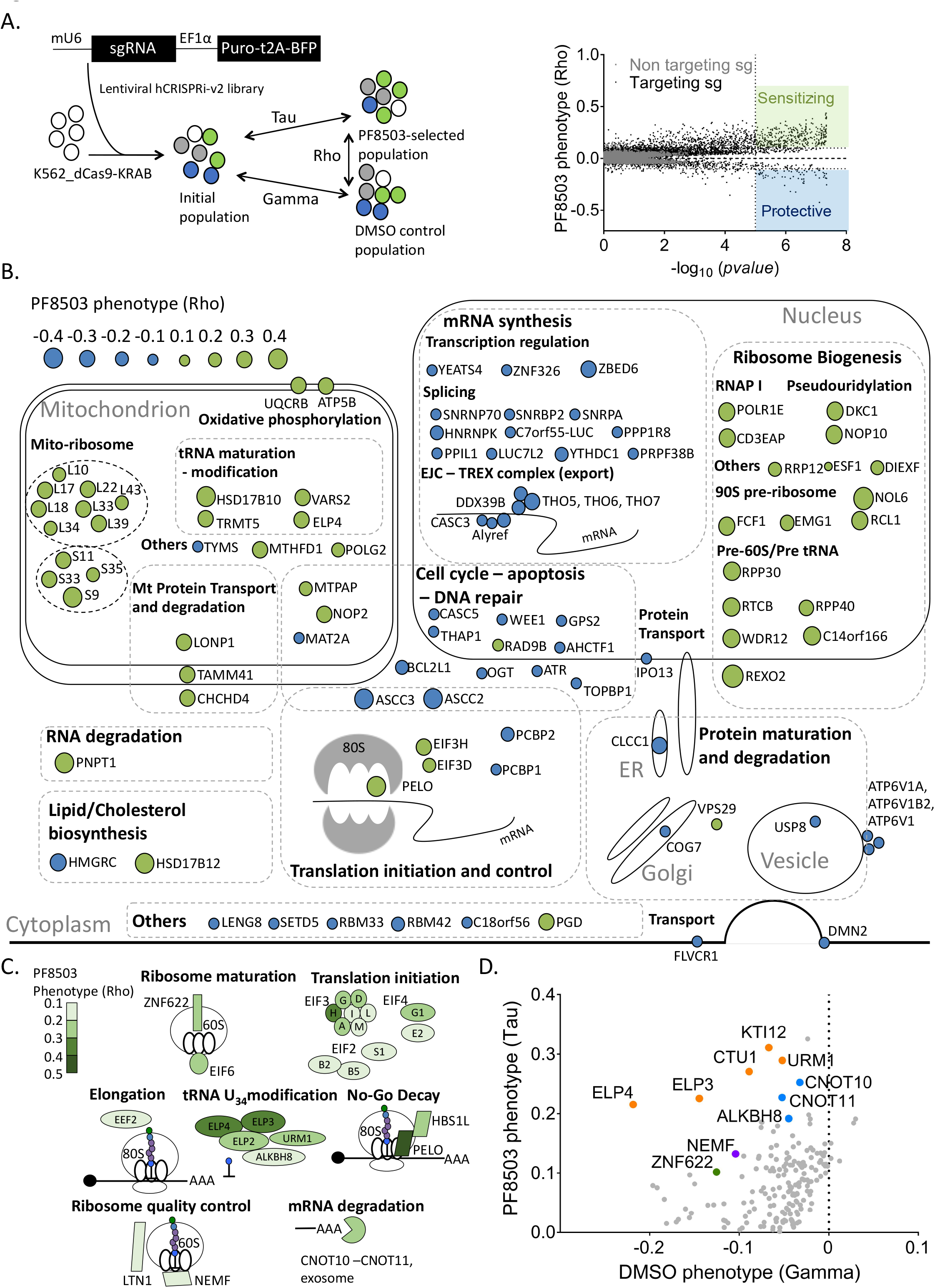
Genome-wide CRISPRi screen to reveal proteins and pathways impacting PF8503 toxicity in K562 cells. (A) Overview of the CRISPRi experiment, carried out using K562 cell lines expressing dCas-KRAB and the hCRISPRi-v2 library. Promoters in the lentiviruses for sgRNA (murine U6, mU6) and Puromycin resistance cassette (EF1α) are shown [17]. After the cells were cultured for 11 days, with or without treatment with PF8503, the quantified genomically-integrated sgRNAs (day 0 and day 11) were used to calculate the effect of each sgRNA on growth without drug (Gamma), with drug (Tau) and the effect of drug treatment (toxin phenotype, Rho). Right, genes with enriched or depleted sgRNAs sensitize (green) or protect (blue) cell viability with respect to Rho, respectively, i.e. with PF8503 treatment. (B) Position of the 50 top protective (blue) and sensitizing (green) genes mapped to cellular pathways. The size of the circles are proportional to the PF8503 Rho phenotype shown in (A). (C) Genes known to be involved in translation which sensitize (green) or protect (blue) cells from PF8503 treatment. (D) Genes with opposite effects in the tau and gamma phenotypes. Some of the proteins of interest involved in tRNA maturation (orange), translation quality control (purple), mRNA degradation (blue) or ribosome assembly (green) are highlighted. In panels (B-D), all phenotypes are the average of the 3 best sgRNAs from experiments carried out in biological duplicate.

Genes from the PF8503 CRISPRi screen were filtered based on their compound-specific phenotypes (Rho > |0.1|) and Mann-Whitney adjusted *p*-value (<0.000001) leading to a total of 452 genes impacting cell fitness (Fig 2A). These genes were then assessed for pathway enrichment using the STRING database [19]. This analysis revealed a clear distinction between pathways protecting or sensitizing the cell to PF8503 toxicity (Fig 2B and S5 Fig). For example, proteins whose expression protects cells from PF8503 toxicity are highly enriched in mRNA synthesis and export pathways (transcription regulation, splicing, EJC-TREX complex) and cell cycle, apoptosis and DNA repair pathways (Fig 2B). By contrast, pathways sensitizing cells to PF8503 are concentrated in the mitochondrion, ribosome biogenesis and translation (Fig 2B and 2C). A third category of proteins are protective in the DMSO control (Gamma) but sensitizing in the PF8503-treated samples (Tau) (Fig 2D), including proteins involved in tRNA wobble nucleotide U_34_ modification [20], 60S ribosomal subunit maturation (ZNF622) and mRNA turnover (CNOT10, CNOT11).

Notably, proteins known to be involved in rescuing stalled ribosomes also affect cell viability in the presence of PF8503, suggesting that ribosome quality control pathways are triggered upon compound-induced stalling (Fig 2B and 2C). No-Go Decay proteins (NGD) Pelota (PELO) and HBS1L, involved in recognizing stalled ribosomes, enhance PF8503-induced cell toxicity, with PELO showing the highest sensitizing phenotype among these hits (Rho = 0.43). Other proteins that sensitize cells to PF8503 include Ribosome Quality Control (RQC) proteins NEMF and LTN1, involved in recycling the 60S subunit of the ribosome, and CNOT proteins involved in mRNA degradation (Fig 2). The sgRNAs most potent in decreasing cell fitness target two of the three subunits that constitute the activating-signal co-integrator complex (ASC-1), subunits ASCC2 and ASCC3. ASCC3 has been implicated in ribosome quality control, by aiding in resolving stalled ribosomes on poly-A sequences [21].

### CRISPRi validation

To further explore the connection between ribosome quality control pathways and PF8503-induced cell toxicity, we first validated the effects of the genes identified in the CRISPRi screen. We constructed CRISPRi K562_dCas9-KRAB cell lines expressing the most active sgRNAs from the CRISPRi screen and tested the effect of PF8503 on these cells using a similar treatment protocol for 7 days (S6A Fig). Knockdown efficiency was confirmed by RT-qPCR and Western blot analysis (**S6B** and S6C Fig). The individual knockdowns were highly correlated with the Rho phenotypes in the CRISPRi screen (Fig 3A, R^2^~0.97), allowing us to use the individual CRISPRi cell lines to evaluate the effects of these proteins on cell fitness.

**Fig 3.**
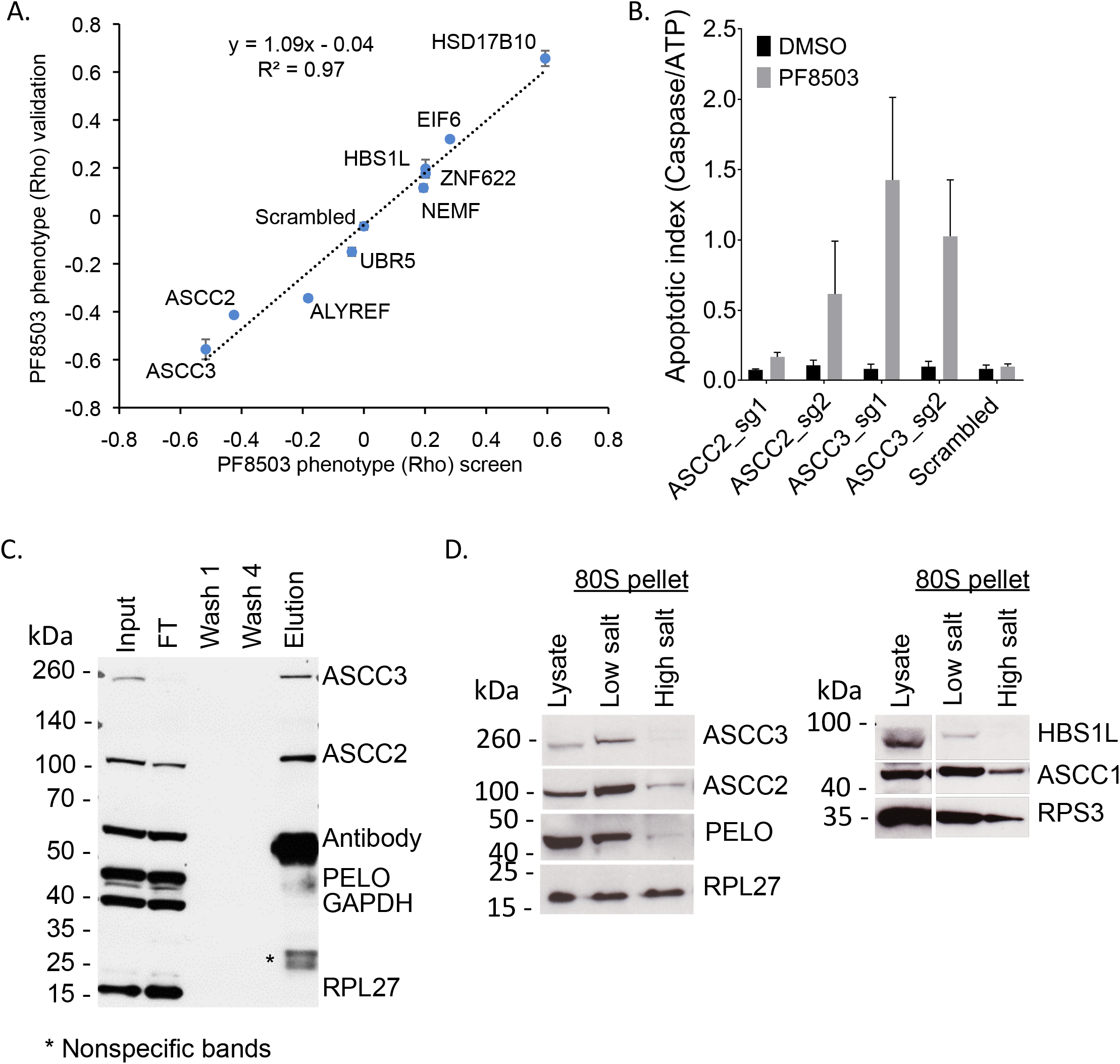
Roles of ASCC2 and ASCC3 in PF8503-dependent toxicty. (A) Comparison of the Rho phenotype in the CRISPRi screen to relative cell viability in individual gene knockouts. K562_dCas9-KRAB cells with individual sgRNAs targeting genes of interest were competed against cells with a scrambled sgRNA, in the presence of 7.5 μM PF8503. Experiments were carried out in biological triplicate with the average log_2_(fold change) and standard deviation shown. (B) Effect of treatment with PF8503 (7.5 μM) on K562-dCas-KRAB cell lines expressing two different sgRNA targeting either ASCC2 or ASCC3 expression. The apoptotic index is the ratio of Caspase 3/7 levels to ATP levels, measured after 6 days of 7.5 μM PF8503 or DMSO control treatment. Experiment performed in triplicate, with the average and standard deviations shown. (C) Western blot of immunoprecipitation of ASCC3 from the cytoplasm of HEK293T cells. Input, cell lysate; FT, flow-through supernatant; Wash 1 and 4, Bead washes; Elution, Proteins extracted from beads. Gel is representative of duplicate experiments. (D) Western blots obtained after isolation of 80S ribosomes from K562-dCas9-KRAB cell lines expressing scrambled sgRNA, using a sucrose cushion at low (200 mM) or high (400 mM) potassium acetate concentration. Gels are representative of experiments carried out in duplicate.

### ASCC2 and ASCC3 impact caspase induction, and interact with cytoplasmic ribosomes

The observation that knockdown of ASCC2 or ASCC3 is highly toxic in the presence of PF8503 suggests a major role of the ASC-1 complex in cell survival upon PF8503 induced translational stress. Induction of apoptosis has been observed in conditions of proteotoxic stress, i.e. downstream of the integrated stress response [22]. We therefore assessed the impact of decreased ASCC2 or ASCC3 expression in the presence of PF8503 by testing for activation of executioner caspases 3 and/or 7 upon long term compound treatment (Fig 3B). Whereas cells transfected with a non-targeting scrambled sgRNA and treated with PF8503 showed no significant induction of caspase 3/7 activation (apoptotic index ~0.1), knockdown of ASCC3 increased the apoptotic index to greater than 1 in the presence of PF8503 but not in DMSO-treated controls. ASCC2 knockdown also elicited a higher apoptotic index in the presence of PF8503 (Fig 3B). By contrast, in a survey of genes identified by the CRISPRi screen, no other knockdown cell lines showed an appreciable induction of apoptosis (S7 Fig).

Although ASCC2 and ASCC3 contribute to cell viability in the presence of PF8503, their role in rescuing stalled translation complexes of the type generated by PF8503 is not established [21]. Furthermore, these proteins in the ASC-1 complex have known roles in the nucleus as transcriptional activators and in the alkylated DNA damage response [23,24]. If these proteins have a direct role in the recognition or rescue of stalled ribosomes, they should interact in the cytoplasm. Immunoprecipitation of ASCC3 in the cytoplasmic fraction of HEK293T cells showed that ASCC2 and ASCC3 interact in the cytoplasm, independently of PF8503 treatment (Fig 3C), although the presence of ASCC2 in the non-bound fraction indicates that not all of ASCC2 is associated with ASCC3. None of three ribosomal proteins tested (RPL27 or RPS3, RPS19) was found in these immunoprecipitations (Fig 3C and S8 Fig). Interestingly, ASCC2 and ASCC3 can be found to co-fractionate with ribosomes in a sucrose cushion of K562 cellular extracts (Fig 3D). In high-salt sucrose cushions, ASCC2 remains bound to the ribosome, whereas ASCC3 levels are greatly reduced, suggesting that the interaction of ASCC3 with the ribosome is indirect and/or unstable. The third member of the ASC-1 complex, ASCC1, also fractionates with the 80S ribosome (Fig 3D), in both low- and high-salt conditions.

### ASCC2, ASCC3 and HBS1L knockdowns do not impact general translation or the ability of PF8503 to induce PCSK9 stalling

Since HBS1L is involved in NGD pathways, and ASCC2 and ASCC3 bind the ribosome and could also contribute to ribosome quality control, we wondered whether their knockdown would impact translation in the presence of PF8503. Using metabolic labelling with the methionine homologue L-AHA, we measured global translation in the presence and absence of PF8503. PF8503 did not decrease global translation in control cells with a scrambled sgRNA, whereas translation was significantly inhibited with non-specific translation inhibitor cycloheximide (Fig 4A). Furthermore, none of the knockdowns of HBS1L, ASCC2, or ASCC3 affected global translation compared to the control cell line, either in the presence or absence of PF8503 (Fig 4B). The ability of PF8503 to induce selective stalling was also unaffected by the knockdowns of HBS1L, ASCC2, or ASCC3. Using a reporter mRNA with codons 1-35 of PCSK9 encoded at the N-terminus of Renilla luciferase, the IC50 for PF8503 was found to be ~0.3 μM for control cells and all three knockdown cell lines (Control, 0.39 +/- 0.02 μM; ASCC2 KD, 0.31 +/- 0.04 μM; ASCC3 KD, 0.26 +/- 0.06 μM; HBS1L KD, 0.39 +/- 0.03 μM; Fig 4D).

**Fig 4.**
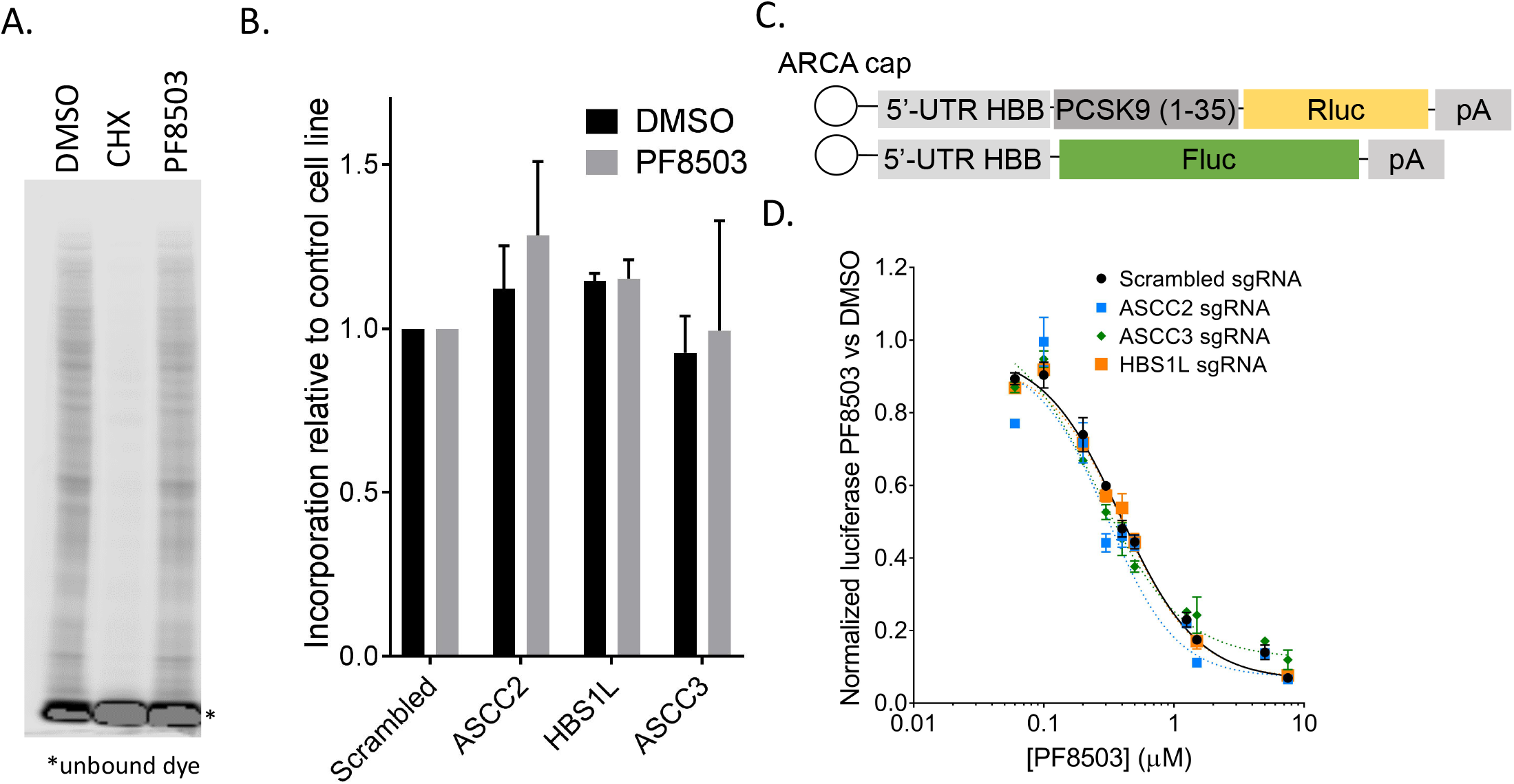
Effect of ASCC2, ASCC3 or HBS1L knockdowns on general translation and PF8503-induced stalling. (A) Metabolic labeling of ongoing translation during 30 min treatment of K562_dCas9-KRAB cells expressing scrambled sgRNA, with PF8503 (7.5 μM), DMSO control, or cycloheximide (100 μg/mL). Shown is IRDye800 labelled L-AHA incorporated into newly synthesized proteins of a representative experiment carried out in duplicate. (B) L-AHA incorporation in newly synthesized proteins in K562_dCas9-KRAB cells expressing scrambled, ASCC2, HBS1L, or ASCC3 sgRNA during 30 min treatment with DMSO or PF8503 (7.5 μM). Ratio of L-AHA incorporation for each knock-down relative to the control cell line are normalized to total protein ratio, determined by Bradford assay. Experiments were carried out in duplicate with the mean and standard deviation shown. (C) Reporter mRNAs used for cell-based assays. ARCA, m^7^G cap with m^7^G nucleotide 3’-*O*-methylated. 5’-UTR HBB, 5’-untranslated region of the *HBB* gene. PCSK9(1-35), codons 1-35 of the *PCSK9* gene. (D) Inhibition of the PCSK9(1-35) reporter mRNA in K562-dCas9-KRAB cells expressing scrambled, ASCC2, HBS1L, or ASCC3 sgRNA, after 6-8 hr treatment with DMSO or various concentrations of PF8503. Experiments were carried out in biological triplicate, with mean and standard deviations at each PF8503 concentration shown.

These results indicate that the effects of knocking down HBS1L, ASCC2, and ASCC3 on cell fitness in the presence of PF8503 is not due to generally lower rates of translation, or to a direct role for these proteins in PF8503-induced stalling.

### Genetic interactions between ASCC3 and RQC and NGD pathways

In order to check whether the effects of ASCC2 and ASCC3 on cell fitness during PF8503 treatment are interdependent, we constructed a cell line in which both ASCC2 and ASCC3 were knocked down using dual sgRNAs, as described in previous Perturb-seq experiments [25] (Fig 5A and S9A Fig). We also generated cell lines in which both ASCC3 and NEMF or ASCC3 and HBS1L were knocked down, to check for genetic interactions with the RQC or NGD pathways, as hypothesized by a role of ASCC3 in a RQC-trigger (RQT) complex [21]. In these experiments, we noted that knockdown of ASCC3 in the context of using dual sgRNAs was not as efficient as the case with single sgRNAs (**S9B** and S9C Fig). We therefore generated a dual-sgRNA cell line with a scrambled sgRNA and ASCC3 sgRNA to serve as a control for the double-knockdown cell lines. We then used competitive growth assays as in the CRISPRi validation experiments to determine the effects of combined knockdowns in the presence of PF8503 (S9A Fig).

**Fig 5.**
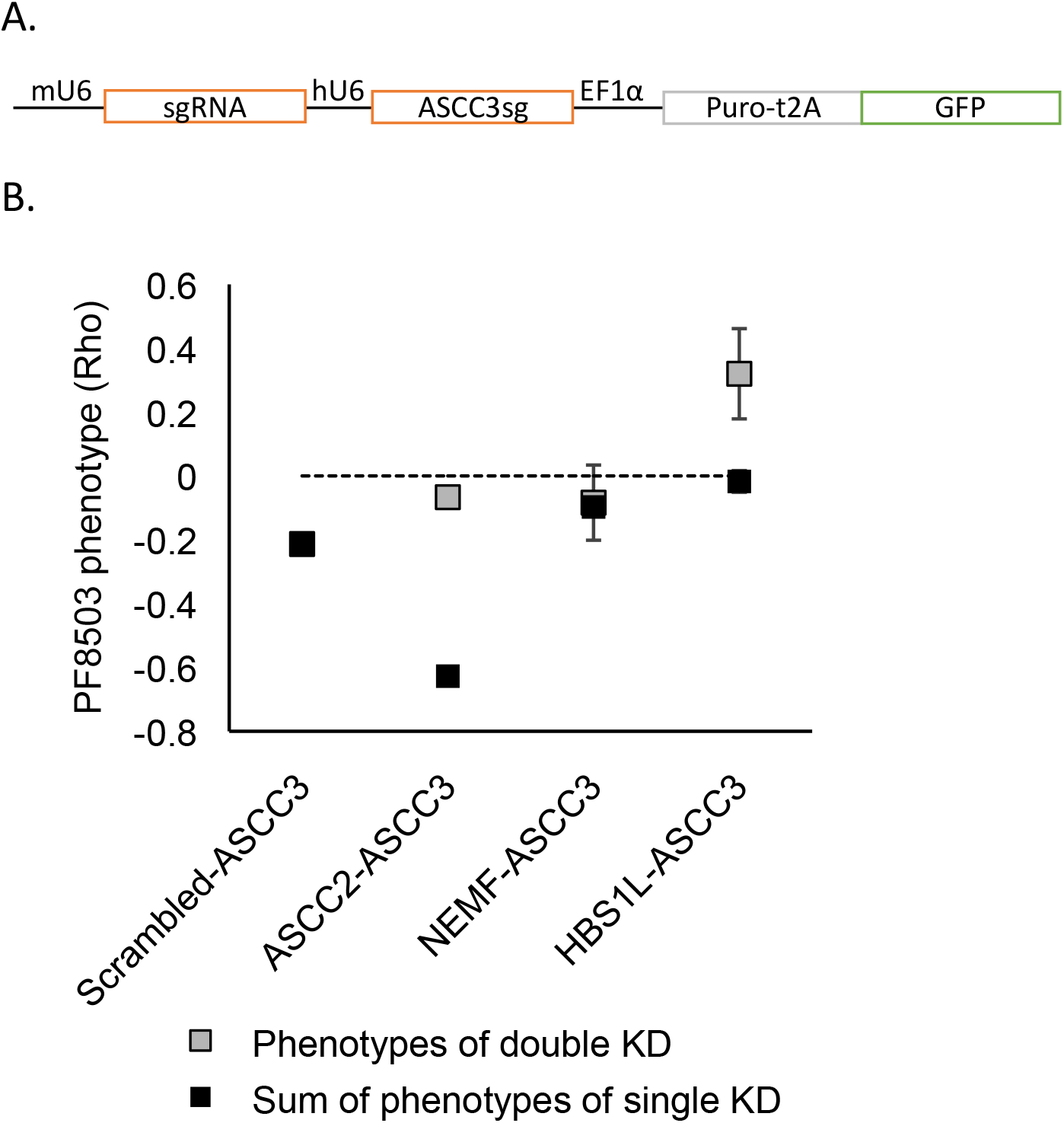
Effects of double knockdowns of ASCC3 and genes involved in translation quality control on PF8503 toxicity. (A) Lentiviral construct used to generate double knockdown cell lines. For each construct, the sgRNA targeting ASCC3 was placed after the human U6 (hU6) promoter, and a second sgRNA (scrambled or targeting ASCC2, NEMF, or HBS1L) was placed after the murine U6 (mU6) promoter. (B) PF8503 phenotype (Rho) obtained in competitive growth assays performed using double knockdown cell lines (grey), compared with the rho phenotype expected from the sum of phenotypes of individual knockdowns. Due to less efficient knockdown of ASCC3 in the dual-sgRNA context, the individual phenotype of ASCC3 knockdown was taken from the cell line expressing mU6-scrambled sgRNA-hU6-ASCC3 sgRNA pair. All other phenotypes were taken from the individual knockdowns in Fig 2A. Experiments were carried out in biological triplicate for ASCC3-NEMF cell line and in 6 replicates for other cell lines, with mean and standard deviation shown.

Interestingly, simultaneous knockdown of both ASCC2 and ASCC3 recovered wild-type fitness in the presence of PF8503 (Fig 5B). This negative epistasis provides strong genetic support that ASCC2 and ASCC3 are part of the same pathway in the context of cell response to PF8503-induced translational stress. On the other hand, the fitness of the NEMF-ASCC3 double knockdown cells is that expected based on simple addition of each protein’s individual phenotype in the presence of PF8503. This does not fully rule out a role of ASCC3 in activating RQC but suggests that these proteins can act in independent pathways. Interestingly, the double knockdown of HBS1L and ASCC3 also showed a negative epistasis, with a phenotype of the double knockdown (Rho~0.3) similar to the phenotype of the single knockdown for HBS1L (Rho~0.2) (Fig 5B). In separate experiments in which lentiviral vectors expressing sgRNAs for HBS1L and either ASCC2 or ASCC3 were introduced into cells sequentially, we also observed a strong negative epistasis (S10 Fig). Importantly, PF8503 retained its ability to stall PCSK9 reporters in the ASCC2-ASCC3 and HBS1L-ASCC3 double knock-down cells (S11 Fig). Taken together, these results suggest that ASCC3 and ASCC2 act in an overlapping pathway with the NGD pathway, which requires HBS1L.

### Comparison of PF8503 with the non-specific translational inhibitor HHT highlights common pathways affected by translation inhibition

In order to ascertain whether the genes identified by the PF8503 CRISPRi screen are specific to selective translational stalling, we compared the PF8503 screen to a CRISPRi screen carried out with the general translation inhibitor HHT, a non-specific translation inhibitor that stalls translation immediately after initiation and early in elongation [26,27][28]. The HHT CRISPRi screen used the same engineered K562_dCas9-KRAB cells, with HHT added at its LD50 value (100 nM). Overall, the HHT-dependent phenotypes for all genes did not correlate well with the PF8503-dependent phenotypes, using the stringent filters applied for the PF8503 CRISPRi screen (Rho_PF8503_ vs. Rho_HHT_, Pearson R=0.11) (Fig 6A). By contrast, the growth phenotypes due to the inherent effects of knockdowns for both CRISPRi screens (Gamma_PF8503_ vs. Gamma_HHT_ and Tau_PF8503_ vs. Tau_HHT_) were well correlated (Pearson R= 0.87 and 0.78, respectively, **S12A** and S12B Fig). In order to compare the PF8503 and HHT effects, we adjusted the *p*-value cutoff for compound-dependent phenotypes with Rho_HHT_ > |0.1| to identify a similar number of genes in the HHT CRISPRi screen (496) when compared to the PF8503 screen (452) (S12 Fig). This allowed us to identify pathways shared between PF8503 and HHT treatments, as well as distinct pathways.

**Fig 6.**
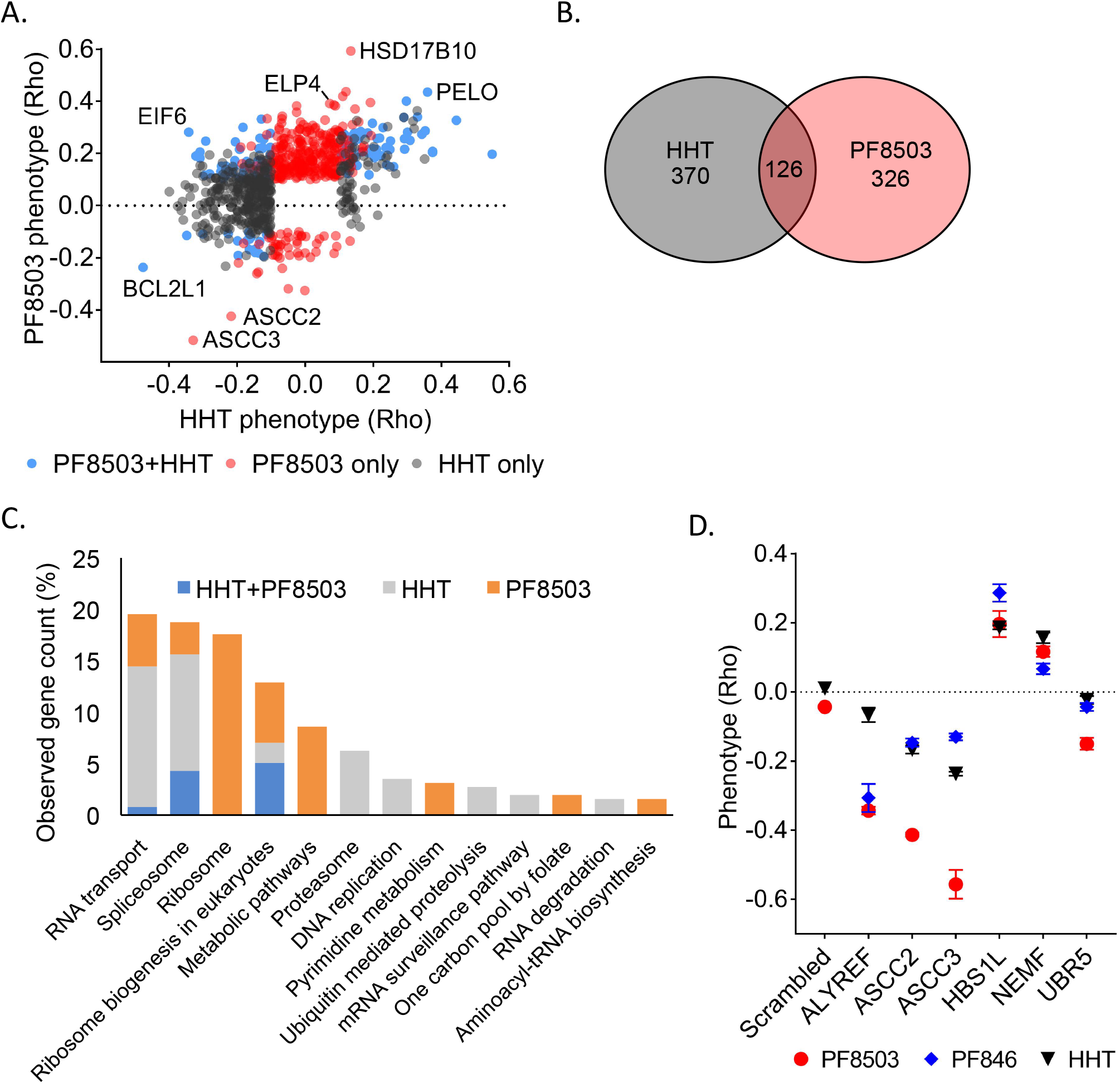
Comparison of PF8503 and homoharringtonine CRISPRi screens. (A) Comparison of gene knockdowns that had significant effects on compound toxicity (Rho phenotype) in CRISPRi screens in the presence of homoharringtonine (HHT) or PF8503. Knockdowns with significant effects in the presence of HHT alone (HHT only, grey), PF8503 alone (PF8503 only, red), or in the presence of either compound (PF8503+HHT, blue) are shown. (B) Venn diagram of significant genes in the HHT and PF8503 CRISPRi screens. (C) Pathways enriched in the common and distinct collection of genes in the HHT and PF8503 CRISPRi screens. Gene count observed corresponds to the number of genes attributed to this pathway by STRING, as a percentage of the total number of genes attributed to a pathway. (D) Phenotypes obtained in K562_dCas9-KRAB competition experiments (scrambled sgRNA as control), with cells treated with three different translation inhibitors: 20 nM HHT, 7.5 μM PF8503, or 7.5 μM PF846. Experiments carried in biological triplicate, with mean and standard deviation shown.

**Fig. 7.**
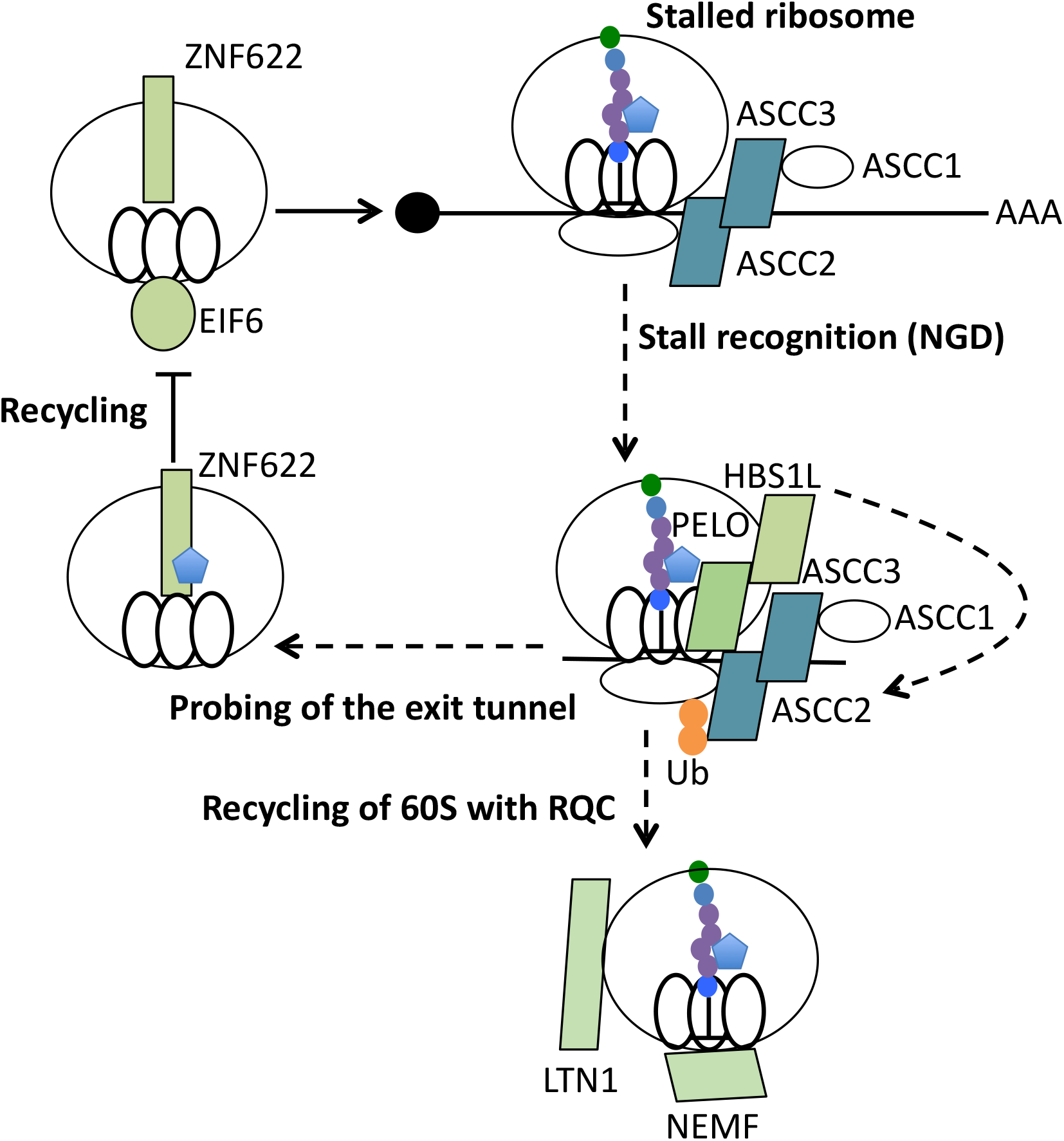
Model for translation quality control pathway that resolves PF8503-induced stalling. PELO/HBS1L detection of PF8503-stalled 80S ribosomes activates a translation quality control pathway that includes ASC-1 (ASCC1-3), which recognizes a K63-linked polyubiquitin signal on the ribosome. These steps are also triggered by HHT. After the 60S subunit is released from the stalled 80S ribosome, RQC may clear the stalled peptidyl-tRNA (LTN1/NEMF). In late steps of recycling, PF8503 bound in the exit tunnel may interfere with quality control functions of ZNF622 and EIF6.

Notably, although only 126 genes were shared between the two screens using the less-stringent cutoff (Fig 6B), a KEGG pathway enrichment analysis performed using the STRING database [19] identified a significant enrichment of the same pathways in the HHT and PF8503 screens, including ribosome biogenesis, the spliceosome and RNA transport (Fig 6C). Proteins with functions related to mitosis, apoptosis, cell cycle, and response to DNA damage also significantly impacted cell fitness in the presence of either of the compounds (**S3 Table** and **S4 Table**). For some of the proteins with shared effects on cell fitness, we compared cell growth in the presence of PF8503, HHT, or PF846 (Fig 6D). Similar effects were observed with all 3 compounds suggesting that these proteins may play a more general role in stress response to translational inhibitors. For example, the two main proteins of the NGD pathway (PELO and HBS1L) and of the RQC pathway (NEMF and LNT1) strongly sensitize cells in the presence of either PF8503 or HHT, suggesting that NGD and RQC could be involved in rescuing stalled or paused ribosomes more generally. By contrast, HHT affected pathways associated with the proteasome and ubiquitin-mediated proteolysis, whereas these did not have a PF8503-dependent phenotype, possibly due to its lower toxicity (**S3 Table** and **S4 Table**). Interestingly, aminoacyl-tRNA biosynthesis pathways were only enriched in the PF8503 CRISPRi screen (Fig 6C). Most of these genes have a Rho phenotype close to 0 in the HHT screen. We also identified that knockdown of two factors involved in late stages of 60S ribosomal subunit assembly, EIF6 and ZNF622, had a positive effect on cell fitness in PF8503-treated cells (Fig 2C and Fig 3A). By contrast, in the presence of HHT, knockdown of EIF6 negatively affected cells, and knockdown ZNF622 did not have a statistically-significant effect (Fig 6A, S12C and S12E Fig).

## DISCUSSION

PF8503 belongs to a new family of compounds able to selectively inhibit the translation of target proteins by the human ribosome. Originally discovered as an orally-available small molecule inhibitor of PCSK9 production [9,10,29], this class of compound could eventually serve as as a new paradigm for designing therapeutics for “undruggable” proteins [2]. These compounds have the unique ability to bind inside the ribosome exit tunnel, allowing them to interact with the protein nascent chain and selectivally stall translation [12]. However, the molecular basis for the strength of compound-induced stalling is still unknown. Furthermore, it is not clear how the cell may respond to ribosomes stalled by these compounds. For example, the cell may clear selectively and non-selectively stalled ribosomes using quality control pathways that remain to be determined. Depending on the strength of the stall, quality control mechanisms could rescue or arrest translation on some transcripts more efficiently than others, participating in the selectivity of these compound and impacting cellular fitness during treatment [30]. Here we used ribosome profiling to map the selectivity landscape of PF8503, allowing comparisons to the related compound PF846 [10], and CRISPRi screens to uncover pathways impacting the toxicity of this class of compound. Taken together, these results identify new connections between ribosome quality control pathways, and should inform future designs of new molecules of this class.

### Selective stalling of translation induced by PF8503

We originally determined the selectivity of compound PF846 (Fig 1A) using human liver-derived Huh-7 cells [10]. Although highly selective, PF846 had some toxic effects in certain cell types and a rat model [10], which would preclude its use for chronic conditions. While optimizing this class of compound, PF8503 (Fig 1A) was shown to have similar potency to PF846 in inhibiting PCSK9 production, as well as a similar cellular toxicity using rat bone marrow as a model for the most sensitive cells to these compounds [9]. Here, we confirmed that PF8503 and PF846 inhibit PCSK9 production by Huh-7 cells with nearly identical IC50 values (Fig 1B), with PF8503 being slightly more toxic when treating human K562 cells (S1 Fig). Although compounds with similar potency to PF8503 and PF846, but with dramatically reduced cellular toxicity, have been identified [9], it is still important to understand why compounds like PF8503 and PF846 decrease cell viability at high doses, as these pathways may arise in the development of new compounds in the future.

Interestingly, although PF8503 and PF846 have similar potency inhibiting PCSK9, they induce ribosome stalling on a surprisingly different array of off-target mRNAs, as determined by ribosome profiling. Of the 46 mRNAs subject to PF8503-induced stalling, only 12 overlap with the mRNAs affected by PF846 (Fig 1C and S3 Fig). For example, translation of *CNPY4*, *TM4SF4*, and *DHFRL1* mRNAs is potently inhibited by PF8503, but these are unaffected or barely affected by PF846 (S3C Fig). PF846 potently inhibits translation of *FAM13B* and *HSD17B11*, whereas these mRNAs are unaffected by PF8503 (S3B Fig). Other mRNAs targeted by one of the compounds may also be stalled by the other, but only at much lower levels that do not lead to a statistically significant reduction in translation past the stall site (**S3 Table** and **S4 Table**, S3 Fig). These results suggest that future efforts to tune the structure of these compounds could lead to selective stalling of new targets beyond PCSK9.

### Genetic interactions with PF8503 identified by CRISPRi

To identify genetic interactions with PF8503 toxicity, we used PF8503 to carry out a whole-genome CRISPRi screen in human K562 cells. By comparing the genetic interactions with PF8503 and the more general translation inhibitor HHT, we were able to identify shared pathways that affect cell viability in the presence of these compounds, as well as ones distinct to each compound class. Although it is possible that common pathways that interact genetically with treatment with PF8503 or HHT (**S3 Table** and **S4 Table**) may reflect a general stress response to translation inhibition, PF8503 does not have an observable effect on global translation (Fig 4A and 4B). This suggests that even the low level of stalled translation induced by PF8503 is sufficient to induce a cellular response that can be detected using CRISPRi. Common pathways that decrease overall translation due a decrease in ribosome biogenesis may lower the overall burden on cells to monitor translation defects, and enable cells to mount a more robust response to PF8503-stalled ribosomes or HHT treatment. An additional explanation for the shared pathways is that HHT, although an inhibitor of late steps of translation initiation [26,27], generates a low level of stalled translation elongation complexes [28] that behave similarly to those generated by PF8503.

Intriguingly, we found that genes involved in tRNA maturation and in particular modification of tRNA nucleotide U_34_ at the wobble position affect cell viability only in the presence of PF8503 (Fig 2C and 2D), not HHT (**S4 Table**). Modifications of U_34_ in tRNAs have been shown to increase the rate of translation [31]. In the present experiments, knockdown of tRNA U_34_ modification enzymes would be predicted to slow down overall translation, providing a protective effect against PF8503-induced stalling similar to that expected from a decrease in ribosome biogenesis. The fact the tRNA synthesis genetically interacts only with PF8503 and not HHT may reflect that fact that PF8503 stalling could be more dependent on the speed of translation, whereas HHT inhibits translation primarily before elongation begins, and would not depend as much on U_34_ modified tRNAs. This interpretation is also consistent with the known binding sites of both drugs. Whereas HHT is bound to the ribosomal A site in the 60S subunit and inhibits the first step of elongation [26,27], PF8503 binds inside the ribosome exit tunnel [12], and likely depends on the speed of translation to stall ribosome nascent chain complexes [10]. Notably, some forms of cancer are dependent on high levels of the enzymes that modify U_34_ in tRNA [32], suggesting that PF8503-class compounds could be developed in the future to target this type of cancer cell addiction.

### Ribosome Quality Control pathways that interact with PF8503

Surprisingly, we also found that knocking down NGD proteins PELO and HBS1L, as well as RQC proteins NEMF and LTN1, had a positive impact on cell fitness upon either PF8503 or HHT treatment (Fig 6A and 6D), suggesting that NGD and RQC are involved in resolving at least some PF8503- or HHT-stalled ribosomes. Notably, whereas the NGD and RQC pathways in yeast have been associated with mRNA turnover, treatment of cells with PF846 has been shown not to lead to decreased levels of PF846-targeted mRNAs [10,11]. The fact that ribosome quality control-related pathways sensitize cells to PF8503 at first seems surprising, since it implies that rescuing translation in response to PF8503-induced translational stalling has a negative impact on cell fitness. However, an alternative explanation is that PF8503-stalled complexes saturate the NGD or RQC pathways in the context of normal cells, preventing these pathways from carrying out their normal quality control functions. Consistent with this saturation hypothesis, the PF846 class of compounds are relatively more toxic in the hematopoietic lineage [9,10], which is more dependent on a functional NGD pathway [33]. Furthermore, ribosome profiling experiments likely miss weak translation stalling events that may occur across the human transcriptome, or those that are robust substrates for ribosome quality control pathways and are therefore not detected in the ribosome profiling analysis. It is notable that some derivatives of the PCSK9-stalling compounds can retain potency without the associated toxicity in rat bone marrow cells [9], suggesting that the NGD- and RQC-dependent cellular responses can be managed for new therapeutic targets.

### Role of ASCC2 and ASCC3 in ribosome quality control pathways

Surprisingly, the genes that protect cells the most from the toxic effects of PF8503-dependent stalling encode components ASCC2 and ASCC3 of the transcriptional activating signal cointegrator complex ASC-1 [24]. Recent experiments have also identified ASCC2 and ASCC3 as integral to the cellular response to DNA alkylation damage [23]. ASCC3 is an RNA helicase and has been shown to be the human ortholog of yeast Slh1 (*SLH1* gene) [21,34]. ASCC2 harbors a CUE (coupling of ubiquitin conjugation to ER degradation) domain that binds K63-linked polyubiquitin chains as part of the DNA alkylation damage response [23]. In yeast, there is emerging evidence of a role for Slh1 in RQC [21,34]. Furthermore, K63 polyubiquitination has been found to be essential for certain stalled ribosome quality control pathways [35]. These studies led to the proposal that Slh1 and Cue3, the presumed orthologue of ASCC3 and ASCC2, are components of a new complex called the RQC-trigger complex (RQT), triggered by ribosomal stalling but not necessary for RQC [21]. Notably, we do not observe a phenotype for the ubiquitin E3 ligase ZNF598 (**S3 Table**), which has been found to ubiquitinate the ribosome in certain contexts of ribosome stalling, i.e. on challenging sequences such as CGA codons and poly(A) stretches in yeast [21,34] and humans [30], and in the presence of colliding ribosomes [30]. These observations prompted us to investigate further the function of ASCC2 and ASCC3 in cellular response to PF8503-induced selective stalling.

We were able to confirm a role for ASCC2 and ASCC3 in response to PF8503-induced stalling using a combination of CRISPRi knockdowns, cell-based assays and cell fractionation (**Fig 3** and S6 Fig). We observed decreased cell fitness and increased induction of apoptosis in PF8503-treated cells when either ASCC2 or ASCC3 were knocked down. Although this could be attributed to nuclear signaling events based on the reported functions for the ASC-1 complex [23,24], we found all three components of ASC-1 associated with cytoplasmic ribosomes (Fig 3D), consistent with previous results connecting K63 polyubiquitination [35] and ASCC3 to translation [21], and suggesting that ASCC1 might be the human counterpart of the third protein of the RQT complex, yKR023W (Rqt4) [21]. Additionally, the fact that ASCC2 and ASCC3 co-IP in the cytoplasm (Fig 3B) is consistent with a role for ASC-1 in translation, independent of its role in DNA alkylation repair and transcription activation in the nucleus. Also supporting a role for ASC-1 in translation quality control, both ASCC2 and ASCC3 knockdowns negatively affect cell fitness of HHT-treated cells (Fig 6A and 6D, S12 Fig).

We also observe that EIF6 and ZNF622, two proteins known to be involved in late stages of ribosomal 60S subunit assembly [36], play a role in PF8503-induced stalling (**Fig 2** and S6D Fig). Knockdown of either factor has a positive effect on cell fitness. Intriguingly, both EIF6 and ZNF622 can be affinity-purified with ASCC2 in a K63-linked polyubiquitin chain-dependent manner [23]. These affinity-purification experiments were conducted in the context of defining the DNA alkylation damage response. However, based on their role in PF8503-induced stalling, it is possible that EIF6 and ZNF622 function in translation quality control pathways, acting downstream of ASC-1.

Importantly, we were able to use double-knockdown cell lines in genetic interaction experiments to identify connections between different translation quality control pathways in human cells. We found that the HBS1L phenotype in the presence of PF8503-induced stalling is dominant over the ASCC3 phenotype (Fig 5B), suggesting that HBS1L and ASCC3 function in a common translation quality control pathway, with ASCC3 involved in steps after translation stall recognition. This result implies that the NGD pathway and ASC-1 intersect to resolve at least some stalled ribosome-nascent chain complexes. The model that ASC-1 acts downstream of HBS1L, after the stalling event, is further supported by the fact that that ASCC2 and ASCC3 knockdowns do not impact general translation or the IC50 for PF8503 (Fig 4B and 4C). Interestingly, we found all three ASC-1 subunits bound to the 80S ribosome independent of active translation (Fig 3D), implying that ASC-1 surveillance of translation may be widespread. By contrast to HBS1L and ASCC3, NEMF which is part of the RQC pathway seems to act independently of ASCC3 (Fig 5B), suggesting that PF8503-induced stalling may not be a “classic” RQC substrate like those identified previously, such as poly-lysine stretches [21,30]. Alternatively, the RQC pathway components may not be as limiting in the presence of PF8503, precluding our ability to detect a genetic interaction. Interestingly, we find that a double-knockdown of ASCC2 and ASCC3 rescues the negative effect of single knockdowns of ASCC2 or ASCC3 upon PF8503 treatment (Fig 5B). We suggest that the negative effect of individually knocking down ASCC2 or ASCC3 could result in a partially activated translation quality control pathway that leads to a build-up of stalled ribosomes that cannot be resolved, thereby leading to severe cell stress.

### Conclusion

The genetic interactions we observe between components of the NGD and RQT, but not RQC, pathways in the presence of specific translational stalling by PF8503-related compounds (**Fig 7**) opens new avenues for exploring the mechanisms of translation quality control pathways in humans. Although the exact role of ASC-1 complex on the ribosome is still unclear, the fact that ASCC2 and ASCC3 strongly impact cell fitness in the presence of PF8503 suggest that ASC-1 may play an essential role in either stall recognition, ribosomal degradation or recycling, or stress signalling to the nucleus. Furthermore, it is possible that ribosome assembly factors EIF6 and ZNF622 also play a role in translation quality control pathways (**Fig 7**), an idea that can now be explored in depth. Combined with the new insights into changes in selectivity when comparing PF8503 and PF846, these results provide a foundation for the future development of compounds that selectively stall the translation of diverse protein targets involved in human disease.

## MATERIALS AND METHODS

### Cells

The human chronic myeloid leukemia (CML) cell line K562, as well as a CRISPRi derivative of this cell line constitutively expressing a catalytically inactive Cas9 fused to a KRAB effector domain (dCas9-KRAB)[17] was kindly provided by the Innovative Genomics Institute, UCSF. K562 cell lines and human hepatocellular carcinoma Huh7 cells (ATCC) were cultured in RPMI 1640 medium (Life Technologies, CARLSBAD, CA, USA) supplemented with 0.2 mM L-glutamine (Glutamax®, Life Technologies), 10% FBS (F4135, Sigma-Aldrich, ST LOUIS, MO, USA), and antibiotics (Penicillin/Streptomycin, 0.1 mg/mL, Gibco) unless otherwise stated. HEK293T cells (UC Berkeley Cell culture facility) were maintained in DMEM (Gibco) supplemented with 10% FBS (Seradigm) and antibiotics unless otherwise stated. Cells tested negative for mycoplasma infection before use.

### PCSK9 inhibition (ELISA)

The compounds PF-06446846 (PF846) and PF-06378503 (PF8503) were synthesized as described in [9] and provided by Pfizer. To determine the IC50 value for inhibition of PCSK9 production, 100 μL of PF846 and PF8503 dilutions in DMSO (0.5% final concentration in media) were added to an overnight culture of 3000 Huh7 cells/mL. PCSK9 production was estimated after overnight incubation by the determination of PSCK9 concentration using a solid phase sandwich ELISA (PCSK9 Quantikine ELISA Kit, RD systems).

### Cell viability and caspase induction assays

96-well plates were seeded with 100 μL of K562 cells at 10^5^ cells/mL and treated with different concentrations of translational inhibitor for 72 hr. For longer experiments, cells were diluted in a new plate and treated again with translational inhibitor in order to remain under the maximum cell density (<10^6^ cells/mL). Cell viability was determined by ATP level measurements (CellTiterGlo2.0, Promega) as an indicator of cell titer and metabolic activity. To determine the level of caspase induction, caspase3/7 levels were measured from 50 μL of cell suspension and ATP levels were measured from the remaining volume (Caspase3/7Glo, Promega). The apoptosis index for each well was calculated as the ratio of Caspase and ATP luminescence signals.

### Ribosome profiling

Overnight cultures of 7 × 10^5^ Huh7 cells in 10 cm dishes were treated with 1.5 μM PF846, 1.5 μM PF8503, or 0.5% DMSO control for 1 hr, in biological triplicate, rinsed with phosphate-buffered saline (PBS) containing cycloheximide (100 μg/mL) and triturated in lysis buffer (20 mM Tris-Cl pH 7.4, 150 mM NaCl, 5 mM MgCl_2_, 1 mM DTT, 100 μg/mL cycloheximide, 1% Triton X-100 and 25 U/mL DNAse I, Promega). The lysates were aliquoted, flash frozen and stored at −80°C until used for ribosome footprint library preparation.

Ribosomes and libraries were prepared as previously described [37]. In short, 300 μL of thawed lysate were digested with RNAse I (Invitrogen, AM2294) for 45 min at room temperature, monosomes were collected using a 1 M sucrose cushion and ultracentrifugation (603,000 g, 2 hr, 4°C), and total RNA was purified using miRNAEasy kit (Qiagen, 217004). RNA fragments of 26 to 34 nucleotides (corresponding to ribosome footprints) were size-selected using a denaturing urea gel. The isolated RNA was dephosphorylated using T4 polynucleotide kinase (New England Biolabs, M0201S), and Illumina-compatible polyadenylated linkers were ligated to the footprint RNAs using truncated T4 RNA ligase 2 (200 U, New England Biolabs, M0242S). First-strand cDNA was synthesized using Protoscript II (200 U, New England Biolabs, M0368L) and circularized using Circligase I (100 U, Epicentre, CL4111K). Ribosomal RNA was depleted using custom oligonucleotides [10] attached to myOne streptavidin beads (Invitrogen, 65001). The cDNA libraries were indexed and amplified using 8 to 14 cycles semi-quantitative PCR with high fidelity Phusion polymerase (New England Biolabs, cat. no. M0530S). Ribosome footprint libraries were sequenced at the QB3 Vincent J. Coates Genomics Sequencing Laboratory, UC Berkeley on an Illumina Hiseq 2500 sequencer.

### Ribosome profiling data analysis

De-multiplexed reads were stripped of 3′ adapters with the FASTX toolkit (http://hannonlab.cshl.edu/fastx_toolkit/index.html) and aligned to human ribosomal RNA sequences using Bowtie [38] to remove ribosomal RNA-mapped reads. A reference transcriptome for Bowtie was built using the UCSC/Gencodev24 known coding Canonical Transcripts of the Grch38 human reference genome [39,40]. To avoid ambiguously mapped reads, only the alignments that mapped to one position of the reference transcriptome were selected for further analysis. To map the reads along the transcripts, an mRNA P site-offset was determined depending on the size of the fragment as 14 nt for fragment of 26 nucleotides or less, 15 nt for fragments of 27 to 29 nucleotides, and 16 nt for fragments of 30 nucleotides or more [41]. Codon maps were then generated for the CDS of each transcript. The data was then processed using custom python scripts and R (**Supplemental document 1**) as previously described with some modifications [10] (S2 Fig). In short, a DMax value was calculated for each transcript as the maximum difference between cumulative normalized reads in PF8503- or PF846-treated and untreated (DMSO control) samples. For each sample, a Z-score transformation of the DMax values was calculated using the scale function in R and the DMax positions with a Z-score greater than or equal to 2 were considered putative stalled transcripts. A differential expression analysis was then performed in R using DEseq for transcripts that had more than 30 counts mapping to CDS reads located 3’ of the DMax position, or 10 codons after the start codon for transcripts lacking a clear DMax value. Reads 10 codons before the end of the transcript were also omitted. Transcripts showing a log_2_ fold change and FDR adjusted *p*-value < 0.05 were considered significant.

To generate ribosomal footprint density plots, the number of ribosomal footprints aligning to each codon position was divided by the total number of reads aligning to the protein-coding regions, then multiplied by 100 to yield reads percentage. All read density plots represent average values for 3 biological replicates. For mRNAs affected by treatment with PF846 and/or PF8503, putative pause sites were found using R and defined as the codon position at which the average read density is at least 10 times higher than the median of the positions on the transcript with more than 0 reads. Footprint densities were then analyzed manually to identify the main stall sites.

Illumina sequencing data and processed ribosome footprints have been deposited at the NCBI Gene Omnibus Database under accession number GSE121981.

### Whole genome CRISPRi negative screen

The “Top5” and “Supp5” plasmid sub-pools of the whole-genome human CRISPRi sgRNA library hCRISPRi-v2 [17] were first pooled to obtain the full 10 sgRNA/gene library and then amplified. Amplification was performed by electroporation into Endura electrocompetent cells (Lucigen, 60242) using the manufacturer’s instructions and amplified in 1 L LB broth with 100 μg/mL ampicillin. The transformation efficiency assured coverage of at least 400X the size of the library. Even coverage of the libraries was confirmed by PCR amplification and deep sequencing.

PF8503 and HHT screens were conducted separately. For each screen, the sgRNA libraries were transduced into K562 cells expressing dCas9-KRAB-BFP as previously described [15]. In short, lentiviral vectors were produced in HEK293T cells by transfection of the hCRISPRi-v2 plasmid library along with packaging (pCMVdeltaR8.91) and envelope plasmids (pMD2.G) using TransIT®-293 Transfection Reagent (Mirius, MIR 2704). Virus-containing media was harvested after 72 hours, supplemented with polybrene (8 μg/mL), and applied to 250 × 10^6^ K562_dCas9-KRAB cells. Cells were spun in 6-well plates (2 hr, 200 g, 33°C) to enhance infection efficiency. Cells were then suspended in fresh culture medium. 60% (for PF8503) or 75% (for HHT) of the cells were infected determined using flow cytometry detection of BFP expression at 2 days post-infection. Cells were then treated with puromycin (0.75 ng/mL, Gibco), for 2 days, at which point the population of BFP-expressing cells reached 80-90% (for both screens). Cells were split into two replicates and the screen was started after one day of recovery in puromycin-free media (T0).

The K562_dCas9-KRAB cells expressing the genome-wide sgRNA library were cultivated in 3 L spinner flasks. For the PF8503 screen, cells were treated 3 times with 7.5 μM PF8503 or vehicle (DMSO), on days 0, 4, and 7. For the HHT screen, cells were treated with 100 nM HHT on day 8. In each screen, the total amount of cells was kept above 250 × 10^6^, and cell density between 0.25 × 10^6^ cells/mL and 1 × 10^6^ cells/mL for coverage of at least 1000X the sgRNA library throughout the screen. Cell size, number, and fraction of sgRNA-containing cells were monitored using a Scepter cell counter and by flow cytometry during the screen. Cells were harvested for library sequencing on the initial cell population (T0) and after 11 days (for PF8503 and corresponding vehicle) or 15 days (for HHT and vehicle).

The CRISPRi screen libraries were prepared from the cells as previously described [15]. Briefly, genomic DNA was extracted from about 300 × 10^6^ harvested cells using Nucleospin XL Blood kit (Machery-Nagel, 740950) and digested overnight at 4°C with SbfI-HF restriction enzyme (New England Biolabs, R3642S). Fragments of about 500 bp, corresponding to the sgRNA cassette, were size selected on a large scale 0.8% Agarose gel and purified using a Qiagen gel extraction kit (28706X 4). The libraries were then indexed and amplified by 23 cycles semi-quantitative PCR using HF-Phusion polymerase (New England Biolabs, M0530S). The libraries were multiplexed and sequenced at the QB3 Vincent J. Coates Genomics Sequencing Laboratory, UC Berkeley, or the Center for Advanced Technology, UCSF, on an HiSeq4000 Illumina sequencer using custom primers as previously described [15]. Computational analysis of the CRISPRi screens was carried out as previously described [15], using the pipeline available on github (https://github.com/mhorlbeck/ScreenProcessing), with the hCRISPRi-v2 library tables/alignment indices. Briefly, the quantified genomically-integrated sgRNAs were used to calculate the effect of each sgRNA on growth without drug (Gamma; T0 vs vehicle), with drug (Tau; T0 vs drug), and the effect of drug treatment only (Rho; vehicle vs drug). Gene-level scores in each condition were obtained by averaging the top 3 sgRNAs for each gene (as ranked by absolute value) and by Mann-Whitney *p*-value of all 10 sgRNAs/gene compared to non-targeting control sgRNAs.

### Lentivirus infection for single and double knockdown cell lines

For each gene candidate to be tested for validation of the CRISPRi results, two sgRNA protospacers giving strong phenotypes in the CRISPRi screen were selected (**S5 Table**) and cloned into a pSLQ1371-GFP or -BFP vector using restriction sites BstXI and BlpI as previously described [17]. Lentiviral vectors were produced in HEK293T cells by transfection of the pSLQ1371-GFP or -BFP sgRNA lentiviral expression vectors along with packaging (pCMVdeltaR8.91) and envelope plasmids (pMD2.G) using TransIT®-293 Transfection Reagent (Mirius, MIR 2704). K562_dCas-KRAB cells were then infected with the resulting lentiviruses in 6-well plates, and puromycin selected (1 μg/mL) two days after infection. The MOI was determined by flow cytometry using GFP or BFP fluorescence before puromycin selection. Knockdown of PELO levels was too toxic to obtain stable cell lines. We therefore used HBS1L knock-down cell lines to explore disruption of NGD.

The double knockdown cell lines were obtained in a similar manner, either by carrying out two sequential lentiviral infections with lentiviruses each expressing one sgRNA. Alternatively, we prepare double knock-down cell lines by a single infection of a lentivirus containing two sgRNA expression cassettes. These plasmids were obtained by ligating a synthetic DNA sequence containing the human U6 (hU6) promoter followed by the sgRNA sequence targeting ASCC3 and a different constant region cr2 [25] downstream of a murine U6 (mU6) promoter and sgRNA expression cassette, in the pSLQ1371-GFP or -BFP vector.

### Western Blot

Cell pellets were lysed using an NP40 lysis buffer (50 mM Hepes pH=7.4, 150 mM KCl, 2 mM EDTA, 0.5% NP40, 0.5 mM DTT) with complete protease inhibitor (Sigma-Aldrich, 4693116001). Proteins were separated on 4-12% Bis-Tris gels (NuPage, Invitrogen, NP0322BOX) and transferred to a nitrocellulose membrane (Ultracruz, sc-3718). Membranes were blocked with 5% skim milk and incubated with primary antibodies overnight. The antibodies used in these experiments are given in **S6 Table**. The secondary antibodies used were conjugated with horseradish peroxide and detected with ECL substrate (PerkinElmer, NEL103E001EA). Western blot films were developed (Optimax, PROTEC), and imaged using Image Studio Lite Software (LI-COR Biosciences, Lincoln, NE, USA).

### RT-qPCR

For the quantification of gene expression of knockdown cell lines, as compared to negative control cell lines with a scrambled sgRNA, total RNA was isolated from cells using an RNA Miniprep kit (Zymo Research, R2060) and analyzed using a SYBR Green 1-step reverse transcriptase real time PCR kit (RNA-to-Ct, Applied Biosystems™, 4389986) on Biorad Quantstudio 3 according to the manufacturer’s instructions. Target-specific primers (**S7 Table**) were designed using Primer-BLAST [42] (https://www.ncbi.nlm.nih.gov/tools/primer-blast/) or the Primer3 web tool ([43], http://primer3.ut.ee/). The cycle threshold (Ct) values were determined using ThermoFisher Connect (https://www.thermofisher.com/us/en/home/cloud.html). Levels of each target RNA were determined relative to the housekeeping gene *PPIA* mRNA levels as internal control for each cell line and the percent of inhibition was calculated using the ΔΔCt method.

### Competitive growth assays

For competitive growth assays, in order to simulate the conditions of the screen, cell lines expressing the scrambled sgRNA were mixed with each target cell line expressing an sgRNA of interest to be tested at a ratio of ~1:1. Each cell line also produced either the GFP or the BFP reporter individually, which allowed the determination of the cell proportion of each cell population over time using flow cytometry. Cell populations were treated with PF8503 (7.5 μM), PF846 (7.5 μM), or HHT (20 nM), near the LD50 for each compound. Growth of the population was also determined by cell counting in order to calculate the enrichment phenotypes as was done in the CRISPRi screen.

### ASCC3 Immunoprecipitation

ASCC3 immunoprecipitation was carried out using HEK293T cells. 50-60% confluent HEK293T cells in a 10 cm dish were treated overnight with 0.05% DMSO or 7.5 μM PF8503. Cells were washed 3 times with HBSS buffer (Gibco, 14025092) and detached by scraping in 1 mL HBSS. Cell pellets were collected after centrifugation and resuspended in 200 μL IP lysis buffer (25 mM Tris-HCl pH 7.4, 150 mM KCl, 0.5 mM DTT, 1X Protease inhibitor, Sigma-Aldrich 4693116001, 4 mM EDTA, 0.5% NP40, 100 U/mL RNAsin, Promega N2511), incubated 10 min on ice, and centrifuged at 4°C for 10 min (14000xg) to remove cell debris. Total cell lysates were incubated with 1 μg of ASCC3 antibody (Bethyl laboratories, A304-015A) for 1 hr at 4°C on a rotator. Then 40 μL of cell lysate was stored at −20°C as the input fraction of Western blot gels. The remaining 160 μL was mixed with 50 μL protein G beads that had been washed and resuspended in 50 μL of NT2 buffer (25 mM Tris-HCl pH 7.4, 150 mM NaCl, 1 mM MgCl_2_, 1X protease inhibitor, Sigma-Aldrich 4693116001, 0.05% NP40, and 100U/mL RNAsin, Promega N2511) and the samples were incubated overnight at 4°C on a rotator. Protein G beads were washed 4 times with NT2 buffer. Proteins bound to the beads were eluted by denaturation in 1X NuPage buffer at 95°C for 10 min and protein content of each fraction were analyzed by Western blot, along with the loading control.

### Metabolic labeling using L-AHA incorporation

Actively-growing K562 cells were washed 3 times with warm PBS and suspended in Methionine free RPMI (Gibco, A1451701), supplemented with 10% dialyzed FBS at equal cell concentrations (~0.5 × 10^6^ cells/mL). Cells were distributed in 6-well plates and incubated for 2 hr in a cell culture incubator at 37°C. Then, 7.5 μM PF8503 or 0.5% DMSO was added to the medium immediately before addition of L-AzidoHomoAlanine (L-AHA) at 25 μM final concentration. Cells were incubated 30 min, washed 3 times with PBS (centrifugation at 400g, 5 min, 4°C) and cell pellets were then flash frozen in liquid nitrogen. Cell pellets were incubated 30 min on ice in 200 μL lysis buffer (1%SDS, 50 mM Tris HCl, pH 8, protease inhibitor 1X, Sigma-Aldrich, 4693116001, and 150 U/mL Benzonase nuclease, Sigma, E1014) and vortexed 5 min. Cell lysates were collected after centrifugation of the cell debris at 18000 g for 10 min at 4°C, then incubated 2 hr with 50 μM of IRDye 800CW DBCO (LI-COR, 929-50000), to label L-AHA with fluorescent dye using a copper-independent Click reaction [44]. Excess dye was removed using a desalting column (Zeba™ Spin Desalting Columns, 40K MWCO, 0.5 mL, 87767). Total protein levels were determined using Bradford protein assay (Biorad, 5000006) for each sample. Total proteins were separated by electrophoresis on 12% or 4-12% Bis-Tris gel (NuPage, Invitrogen, MES-SDS buffer, NP0002) and the gels were washed with PBS. Newly synthesized proteins were labelled with IRDye800 and were imaged in the gel on a LI-COR Infrared imager (Odyssey, 800 nm wavelength channel). After imaging of labeled proteins, total proteins were stained with Coomassie (SafeStain Blue, Invitrogen, LC6060). Images were analyzed using LI-COR Image studio LiTe software (LI-COR Biosciences, Lincoln, NE, USA).

### Production of mRNA

All mRNA reporters were produced from plasmids linearized using PmeI (New England Biolabs, R0560S), in which the mRNA was encoded downstream of a T7 RNA polymerase promoter. Reporters used for in-cell assays were transcribed, capped and tailed using HiScribe™ T7 ARCA mRNA Kit (with tailing, New England Biolabs, E2060S) following the manufacturer’s instructions. Reporters used for *in vitro* translation assays and sucrose cushions were transcribed using HiScribe™ T7 Quick yield (New England Biolabs, E2050S). All mRNA reporters were purified by LiCl precipitation (2.5 M final concentration), washed with 70% ethanol, and quantified by spectrometry (NanoDrop® ND-1000 UV-Vis Spectrophotometer).

### Cell extract preparation and *in vitro* translation

K562_dCas-KRAB cell lines expressing different sgRNAs were grown in 3 L spinner flasks and collected when the density was ~0.5 × 10^6^ cells/mL. Cells were pelleted at ~100 g for 4 min at 4°C, washed in isotonic buffer (20 mM MOPS/KOH pH 7.2, 130 mM NaCl, 5 mM KCl, 7.5 mM magnesium acetate, MgOAc, 5 mM sucrose), and suspended in 1 – 1.5 volumes of cold hypotonic buffer (10 mM MOPS/KOH pH7.2, 10 mM KCl, 1.5 mM MgOAc, 2 mM DTT). Cells were incubated in hypotonic buffer for 10 min on ice and shredded 15X through a 26G needle. Cell extracts were cleared by a 4 min centrifugation at 10000 g, 4°C and cell extract aliquots were flash frozen and stored at −80°C until used.

*In vitro* translation reactions (IVT) contained 50% thawed cell extract in energy and salt mix buffer at the final concentrations: 20 mM MOPS/KOH pH 7.5, 0.2 mM ATP, 0.05 mM GTP, 15 mM creatine phosphate, 0.1 mg/mL creatine phosphokinase, 1 mM DTT, 1.5 mM MgOAc, and 200 mM KCl, 1x amino acid mix (88x amino acid mix was prepared with 16.7x MEM essential amino acids, 33.3x MEM nonessential amino acids, and 6.7 mM glutamine, Life Technologies, 11140050, 11140050, and 25030024), 40 ng/μL reporter mRNA, 0.05% DMSO or various concentrations of the specific translation inhibitor. Total reaction volumes of 10 μL to 100 μL were incubated in a thermocycler at 30°C for 15 to 45 min. Nanoluciferase signals were determined in each well by mixing 8 μL of the IVT reaction with 50 μL of luciferase substrate (NanoGlo, Promega, N1110), and measured using a Veritas microplate luminometer (Turner Biosystems).

### Sucrose cushion

To test the effect of salt concentration on protein adsorption to ribosomes in sucrose cushion experiments, potassium acetate (KOAc) was added after the IVT reaction to a final concentration of 400 mM (high salt) or 200 mM (low salt) prior to layering on the sucrose cushion in MLA-130 tubes (Beckman Coulter, 343778). 100 μL of the adjusted IVT were layered on top of a 600 μL sucrose cushion (1 M sucrose, 1 mM DTT, 20 U/mL Superasin, Invitrogen, cat. no. AM2694, 4 mM HEPES 7.4, 30 mM KOAc, 1 mM MgCl_2_) and ultracentrifuged at 603,000 g for 50 min at 4°C (Optima MAX Ultracentrifuge, Beckman Coulter). To identify proteins that co-isolate with the ribosome, the ribosome pellet was resuspended in 1X Nupage sample buffer (NuPage, NP0008) and protein content of this fraction was determined by Western blot.

### Cell-based Luciferase reporter assays

The mRNA reporters used in cell reporter assay contained the 5’-untranslated region (5’-UTR) of β-globin (*HBB*) followed by either the stalling sequence of PCSK9 (codons for amino acids 1-35), the p2A sequence [45] and Renilla luciferase (PCSK9(1-35)-Rluc), or by just the Firefly luciferase sequence (Fluc). Briefly, 125 μL of K562 cells were seeded in 96-well plates at ~0.4 × 10^6^ cells/mL and incubated overnight (37°C, 5% CO_2_). The test reporter (PCSK9(1-35)-Rluc) and the control reporter (Fluc) were co-transfected (0.05 μg per mRNA per well) in K562 cell lines using JetMessenger mRNA transfection kit (Polyplus, 150-01) according to the manufacturer’s instructions. Serial dilutions of 200x PF8503 were prepared in DMSO and added to the cells immediately after transfection (0.5% DMSO final per well). Rluc and Fluc luminescence signals were measured 7-8 hr after transfection with Dual-Glo (Promega, E2920) on a Veritas microplate luminometer (Turner Biosystems). The normalized signal for each well (Rluc/Fluc) was calculated and normalized to the signal of non treated (DMSO only) wells. IC50 values were calculated using GraphPad Prism version 7.00 for Mac (GraphPad Software, La Jolla California USA, www.graphpad.com).

## ACKNOWLEDGMENTS

We thank R. Dullea, S. Liras, K. McClure, W. Li, R. Green, A. Buskirk, C. Wu, and C. Wolberger for helpful discussions. This work was funded by the Pfizer Emerging Science Fund, and by the NIGMS (P50-GM102706 to JC and JSW and U01 CA168370, R01 DA036858 to JSW). The Vincent J. Coates Genomics Sequencing Laboratory at UC Berkeley is supported by NIH S10RR029668, NIH S10OD018174 and NIH S10RR027303 instrumentation Grants. JSW is a Howard Hughes Medical Institute Investigator. LAG is supported by NIH/NCI R00 CA204602 and DP2 CA239597 and the Goldberg-Benioff Endowed Professorship in Prostate Cancer Translational Biology.

## SUPPORTING INFORMATION

**S1 Fig.**
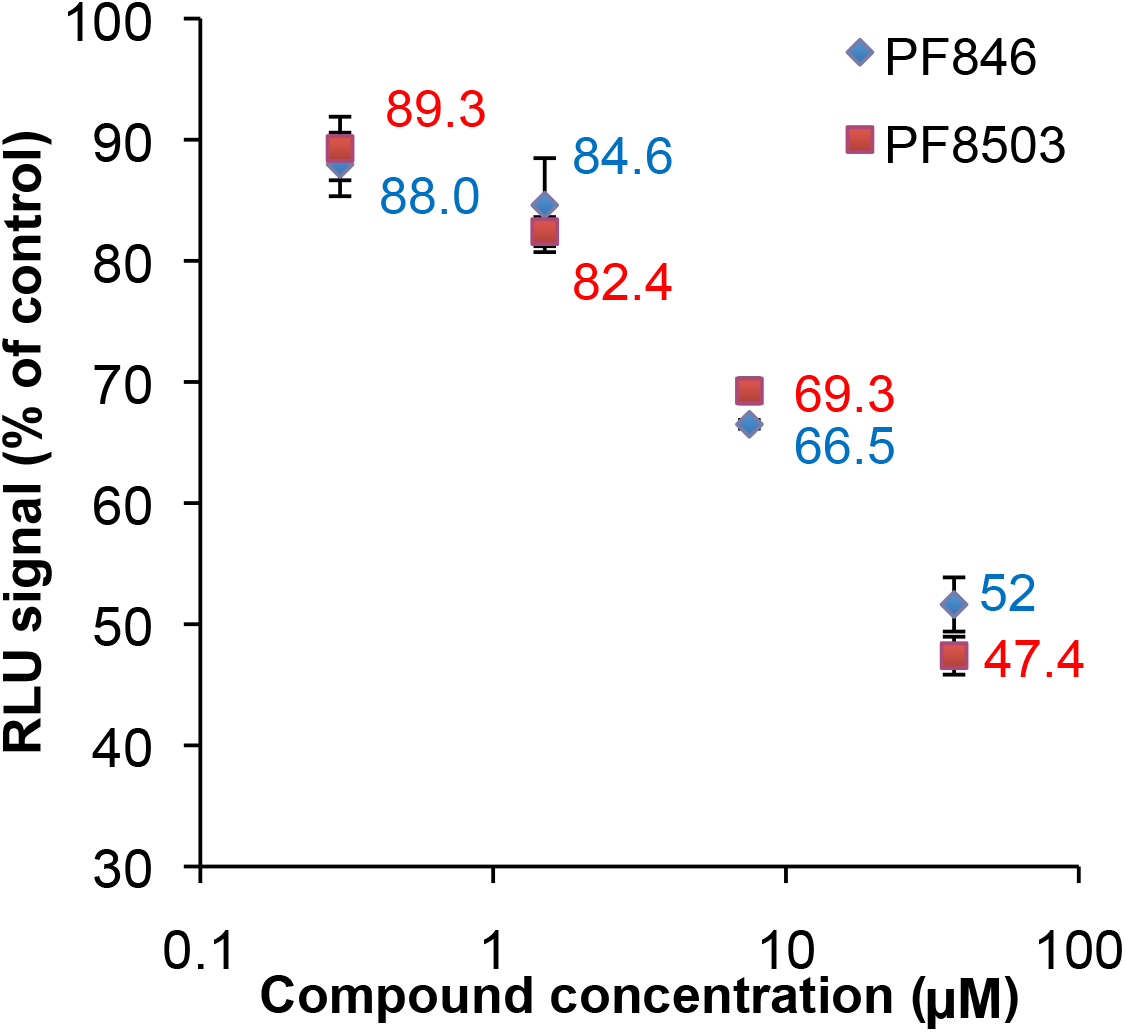
Viability of K562 cells treated with PF846 or PF8503. K562 cells were treated with various concentrations of PF846 or PF8503 in 96-well plates for 48 hr. Viability was measured as percentage of ATP signal in treated vs. not treated (control) wells. Shown are the are the mean and standard deviations of Relative Light Unit (RLU) measurements from 3 replicates each.

**S2 Fig.**
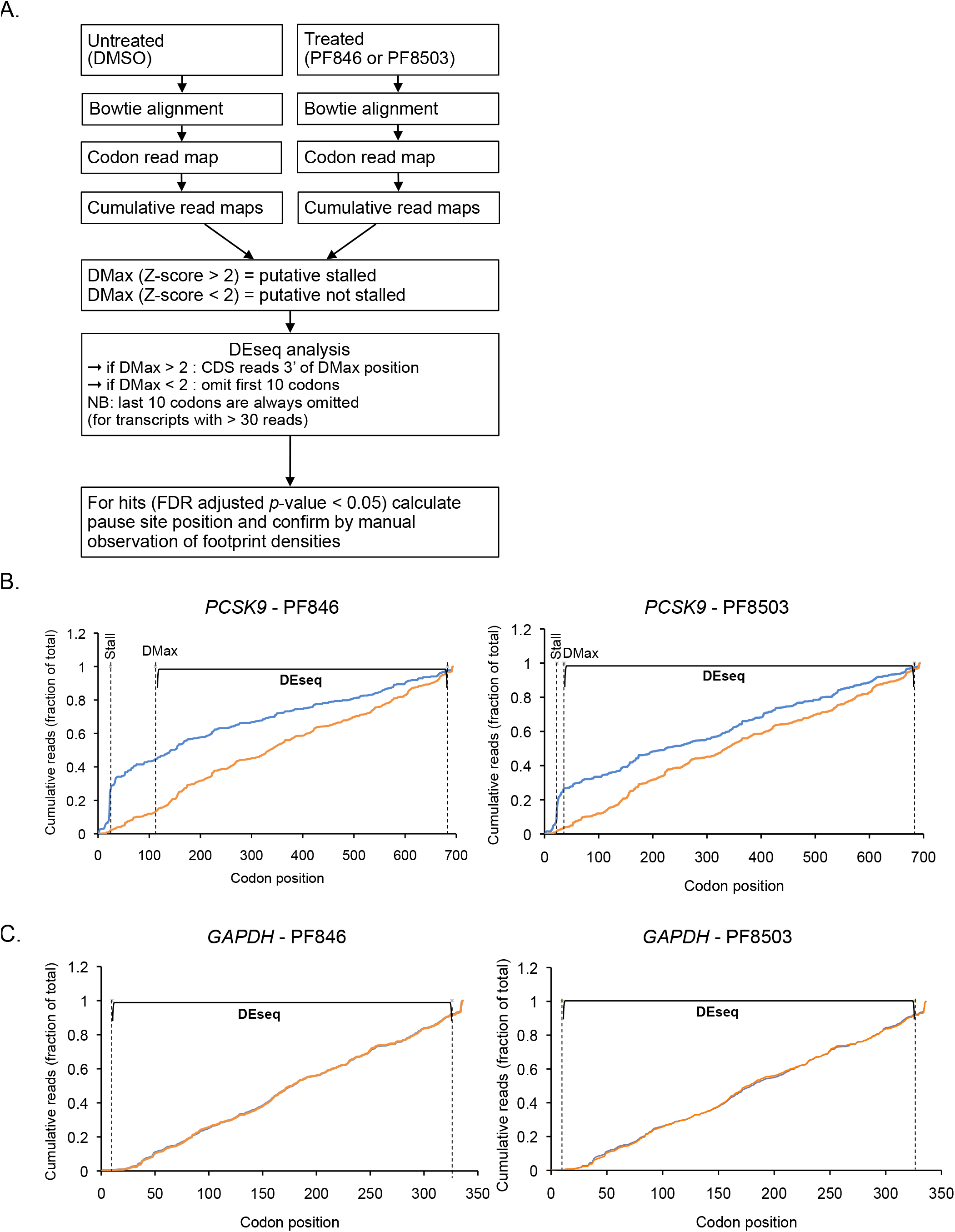
Ribosome profiling data analysis protocol to identify compound-induced stall sites. (A) Data processing flowchart, beginning with ribosome profiling libraries. (B) Examples of compound-induced stalling on *PCSK9* mRNA, showing cumulative read maps, stall site positions, DMax positions, and regions used for differential expression (DEseq) analysis. (C) Example in which compound-induced stalling is not observed (*GAPDH* mRNA). Region used for differential expression analysis (DEseq) is marked.

**S3 Fig.**
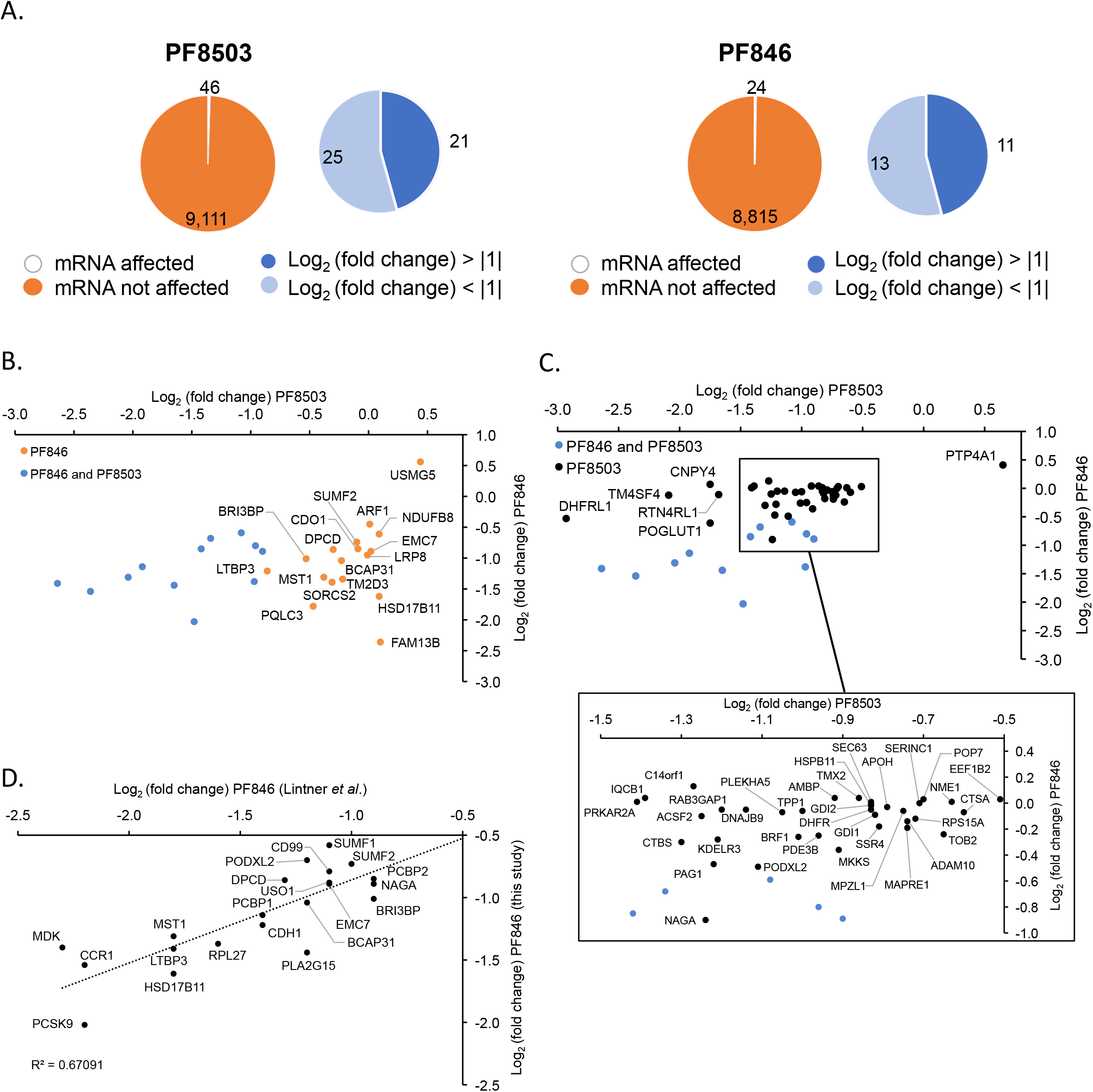
Transcripts affected by PF846 and PF8503 in Huh-7 cells, as revealed by ribosome profiling. (A) Transcripts quantified from Huh-7 cells treated with 1.5 μM PF846 or 1.5 μM PF8503 for 1 hr before harvesting and ribosome protected RNA fragment library preparation. The log_2_(fold change) values correspond to the ratio of reads in compound-treated vs. control cells, summed 3’ of the DMax position, as described in the Materials and Methods and diagrammed in (S2 Fig). Number of mRNAs affected by PF846 or PF8503 (with adjusted *p*-value < 0.05, blue) among the total transcripts that could be analyzed (orange). Data are from three replicates, using the mean values of log_2_(fold change). (B-C) Values of log_2_(fold change) obtained from PF8503- or PF846-treated cells for shared PF8503 and PF846 targets (blue) and targets detected only in the PF846 treatment (orange) or only in the PF8503 treatment (black). Shared targets are also shown in Fig 1C. Data are from three replicates, with the mean values shown. (D) Comparison of the log_2_(fold change) values from PF846-treated cells obtained here with those from PF846-treated Huh-7 cells described previously [10].

**S4 Fig.**
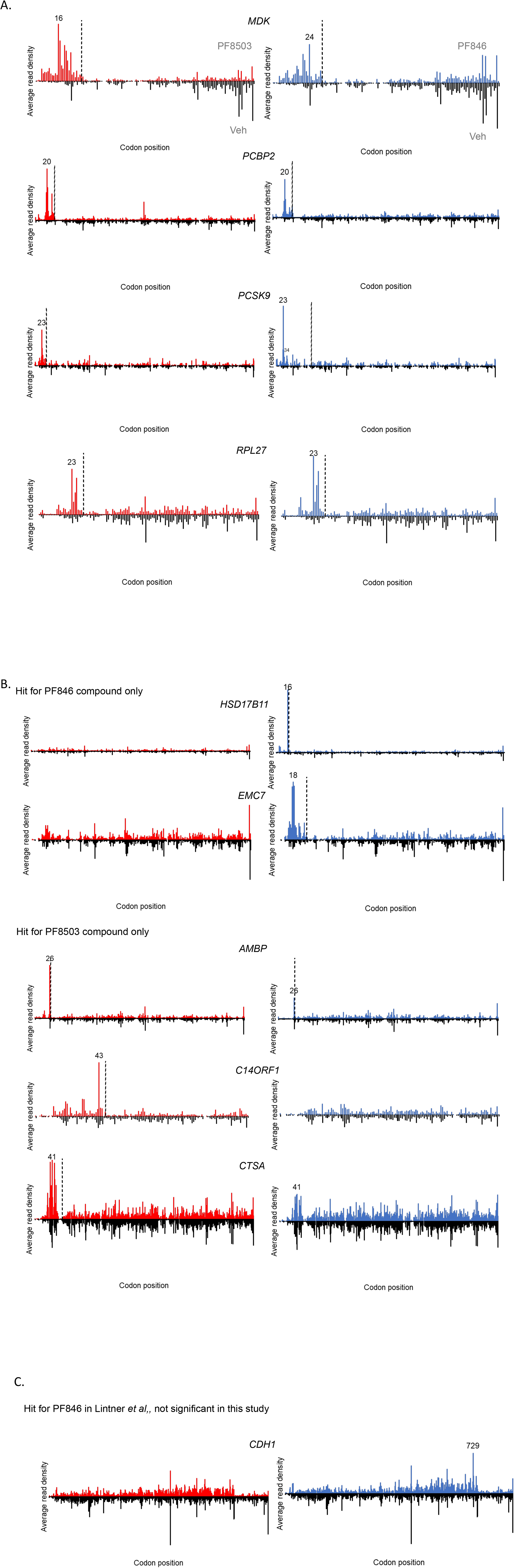
Example ribosome profiling of transcripts affected by PF846 and PF8503 in Huh-7 cells. (A) Examples of transcripts impacted by both PF846 and PF8503. Average ribosome footprint densities along transcripts at codon resolution, from PF8503 (red), PF846 (blue) and Veh (DMSO, black) treated cells. The Veh profiles are shown as mirror images of the compound treatments. The main stall codon is marked for each condition and the dotted line represents the DMax position. (B) Example of transcripts impacted by only one compound. (C) *CDH1* transcript showing a late stall only in the presence of PF846. Note, in the present experiments with PF846, *CDH1* did not pass the DMax Z-score cutoff (**S2 Table**). In panels (A-C), the experiments were carried out in biological triplicate.

**S5 Fig.**
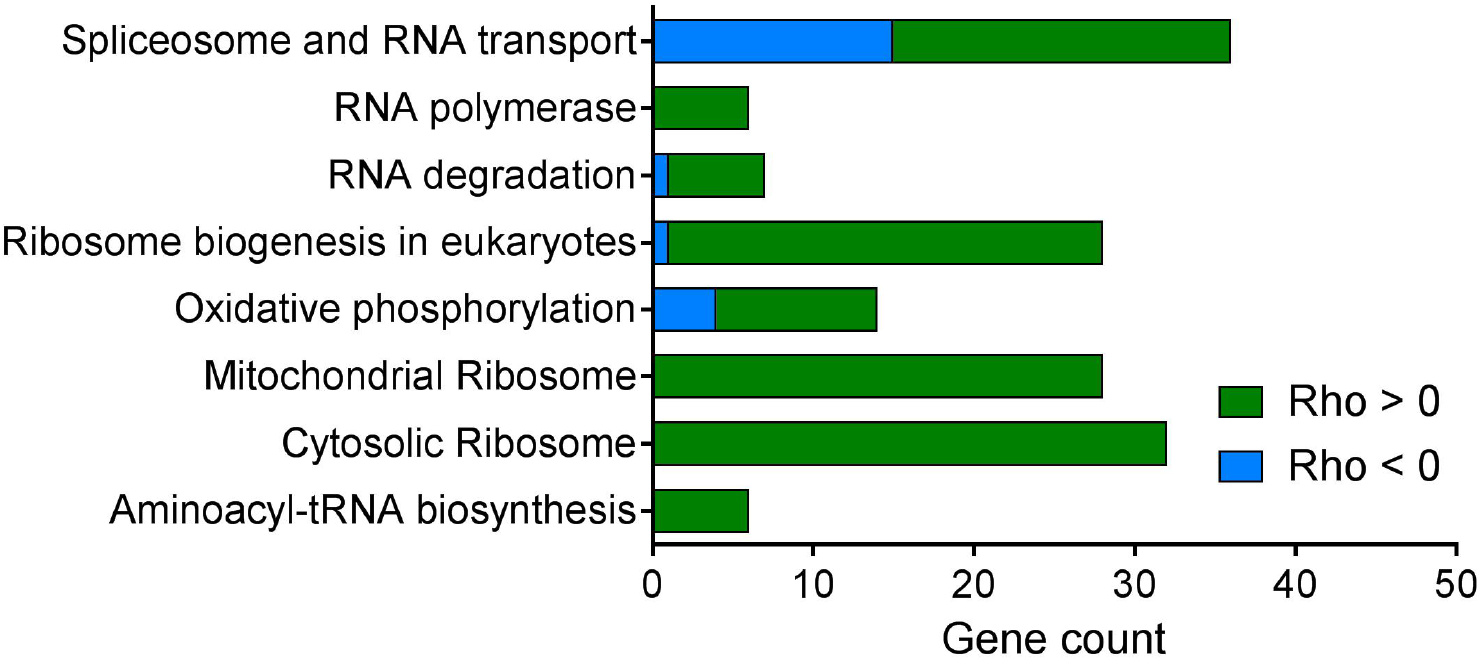
Pathways enriched in the CRISPRi genomic screen of genetic modifiers of PF8503 toxicity. Pathways from STRING database analysis, with genes whose knockdown sensitizes (blue) or protects (green) cells from PF8503 toxicity.

**S6 Fig.**
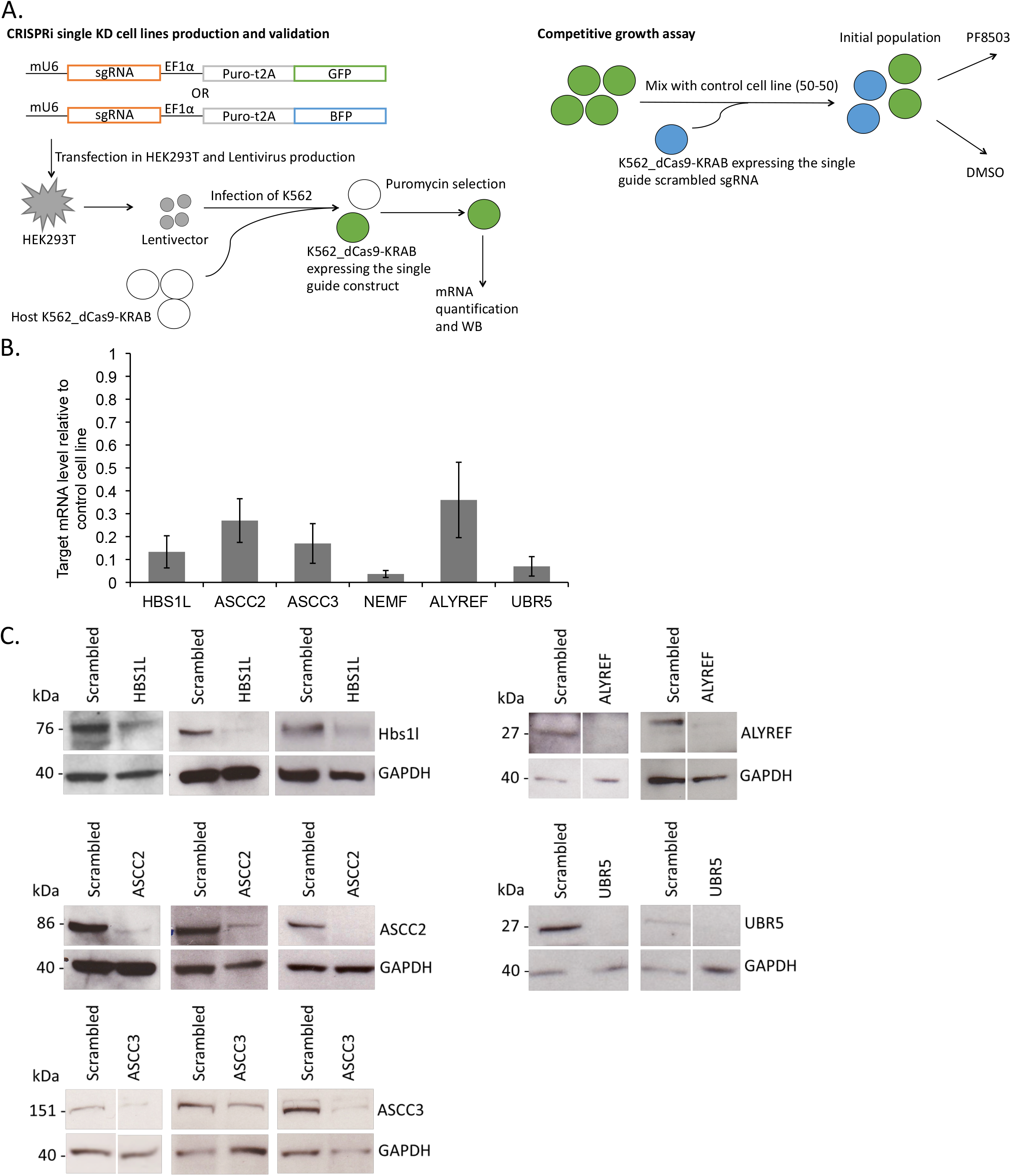
Knockdowns of single-gene expression by individual sgRNAs in K562_dCas9-KRAB cells. (A) Schematic of generation and validation of sgRNA-mediated knockdown in individual cell lines. Lentiviral vectors expressing puromycin resistance and BFP or GFP were used to ensure near-complete lentiviral infection. The resulting cell populations were used for RT-qPCR or Western blot analysis. (B) Levels of mRNAs for targeted genes, as determined by RT-qPCR. Measurements carried out in triplicate, with mean and standard deviation shown. (C) Western blots of proteins whose mRNA transcription was targeted by individual sgRNAs. Each Western blot is from cell lines used for triplicate experiments.

**S7 Fig.**
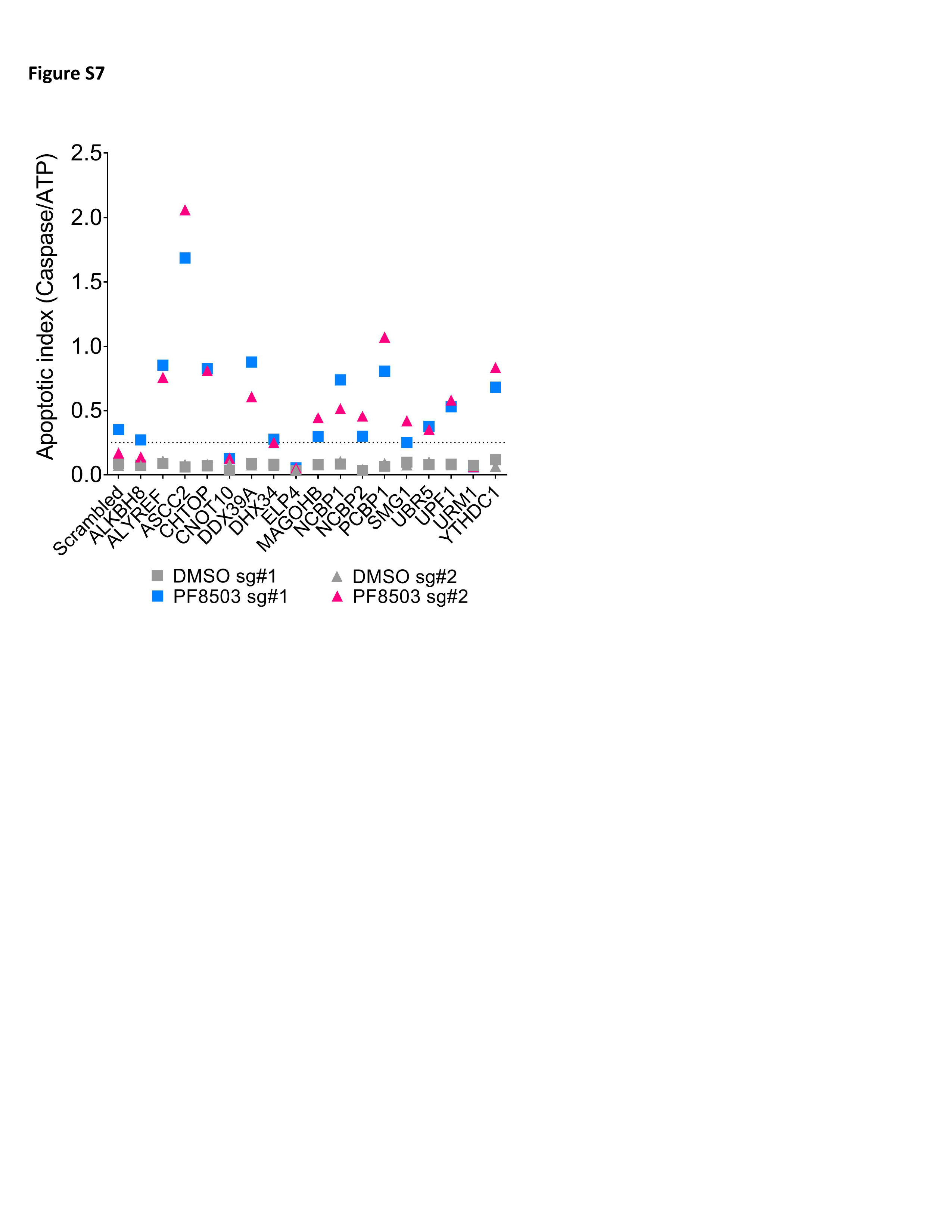
Apoptotic index of individual sgRNA-mediated knockdown cell lines. Survey of the apoptotic index (Caspase 3/7 levels/ATP levels) for cell lines expressing either of two different sgRNA targeting select proteins identified from the CRISPRi screen. Cells were incubated with 7.5 μM PF8503 for 6 days.

**S8 Fig.**
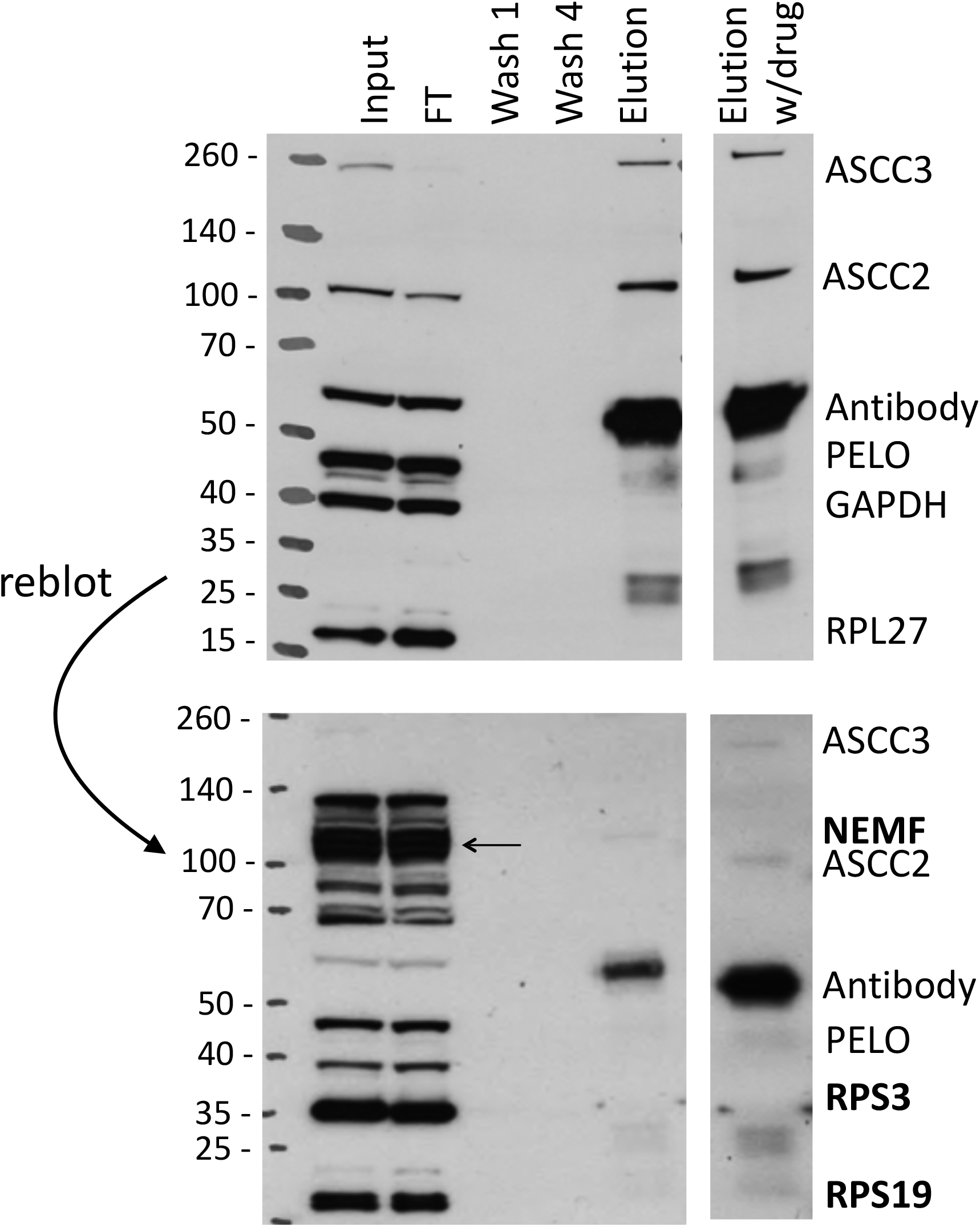
Western blots of ASCC3 immunoprecipitation. Full Western blot gels shown in Fig 3C. Top, blotted with antibodies against ASCC3, ASCC2, PELO, GAPDH, and RPL27. Bottom, membrane stripped and re-blotted for NEMF, RPS3, and RPS19 (bold). NEMF position is indicated by an arrow.

**S9 Fig.**
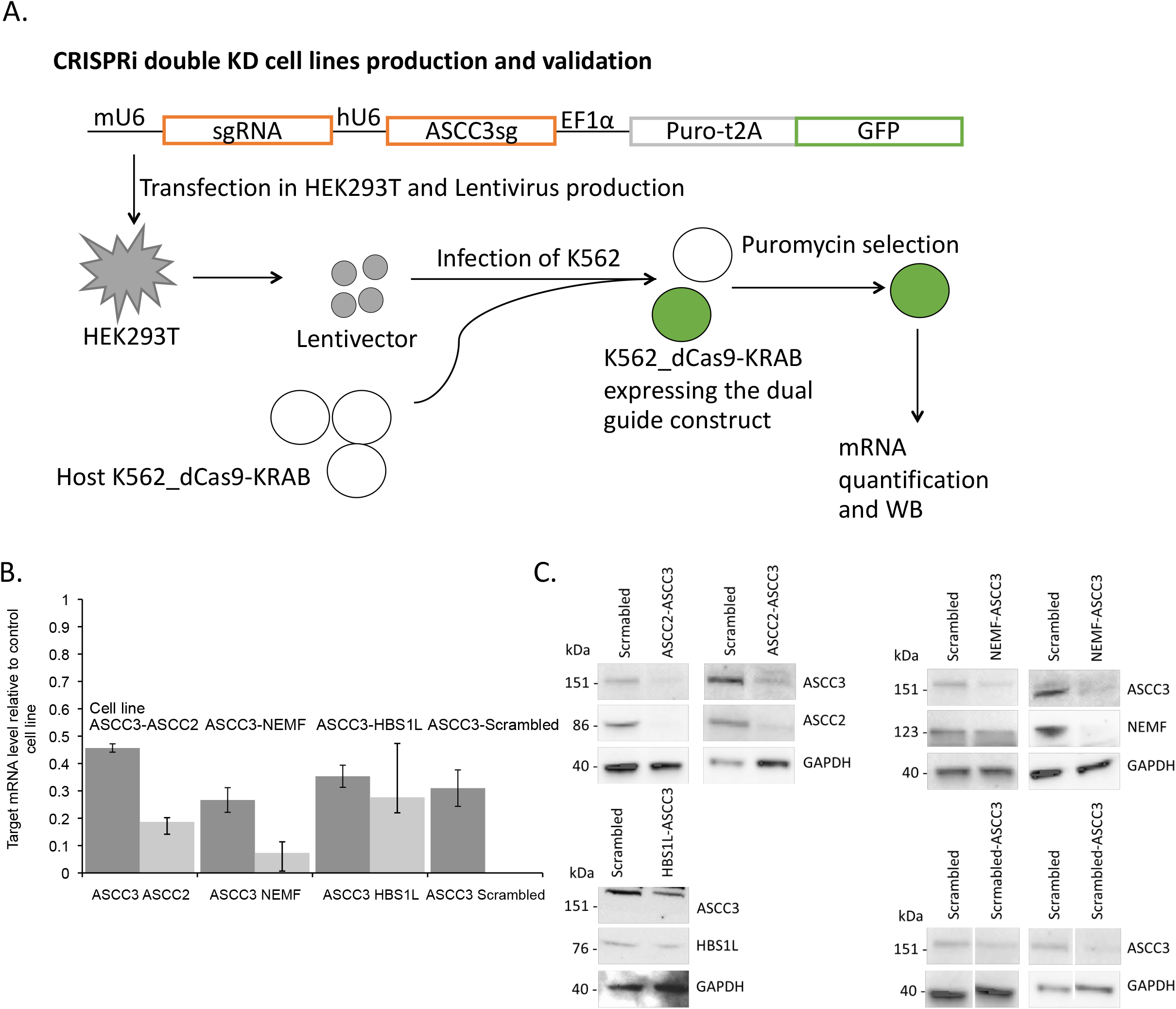
Generation of double knockdown cell lines using dual sgRNAs in K562_dCas9-KRAB cells. (A) Schematic of the construction of double knockdown cell lines. ASCC3 sgRNA expressed from the human U6 (hU6) promoter; second sgRNA expressed from the murine U6 (mU6) promoter. Puromycin resistance (Puro) and GFP expression were used to enrich lentivirally infected cells. The mRNA levels were determined using RT-qPCR, normalized to the housekeeping gene *PPIA* mRNA levels. (B) Target mRNA levels in double knockdown K562 cell lines expressing dCas9-KRAB and HBS1L, ASCC2, or NEMF sgRNAs along with ASCC3 sgRNA. Experiments carried out in triplicate. (C) Western blot analysis of corresponding protein levels in double knockdown cell lines, compared with cells expressing a scrambled guide RNA (NC, negative control). Blots were made using lysates from cells lines grown in triplicate.

**S10 Fig.**
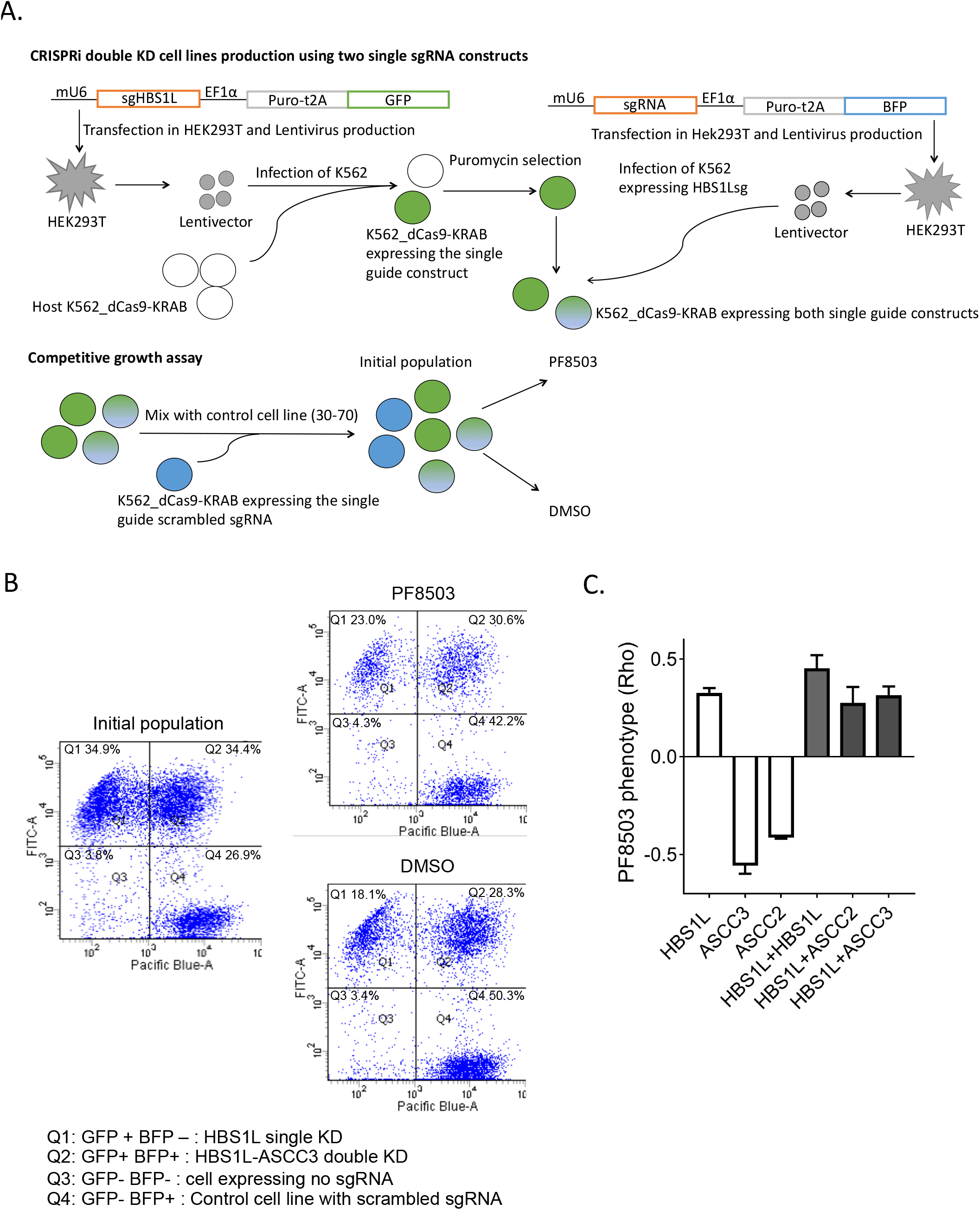
Double knockdown cell lines using sequential transfection. (A) Strategy used to generate double knockdown cell lines. Lentiviral vectors expressing single sgRNAs were used in serial infections to generate double-knockdown cells. Cells expressing sgRNA targeting *HBS1L* (HBS1L sg#2) with a GFP reporter were first validated for HBS1L mRNA knockdown and HBS1L protein knockdown (S6 Fig). These cells were then retransfected with a second lentivirus expressing an sgRNA targeting *ASCC3*, *ASCC2*, or *HBS1L* (HBS1L sg#1), with a BFP reporter. Populations of cells after Puromycin selection could then be scored for both GFP or BFP expression to indicate dual infection with the two lentiviruses. (B) Example FACS analysis of HBS1L-ASCC3 double-knockdown cells before and after selection in the absence or presence of 7.5 μM PF8503. (C) PF8503 toxicity phenotype (Rho) obtained from competitive growth assays in the presence of 7.5 μM PF8503 and scored using FACS analysis of GFP and BFP expressing cells as previously described [15,17]. Individual knockdown cell lines (open bars) and double knockdown cell lines (filled bars) are from experiments carried out in 2 replicates, from two independent transfections with mean and standard deviation shown.

**S11 Fig.**
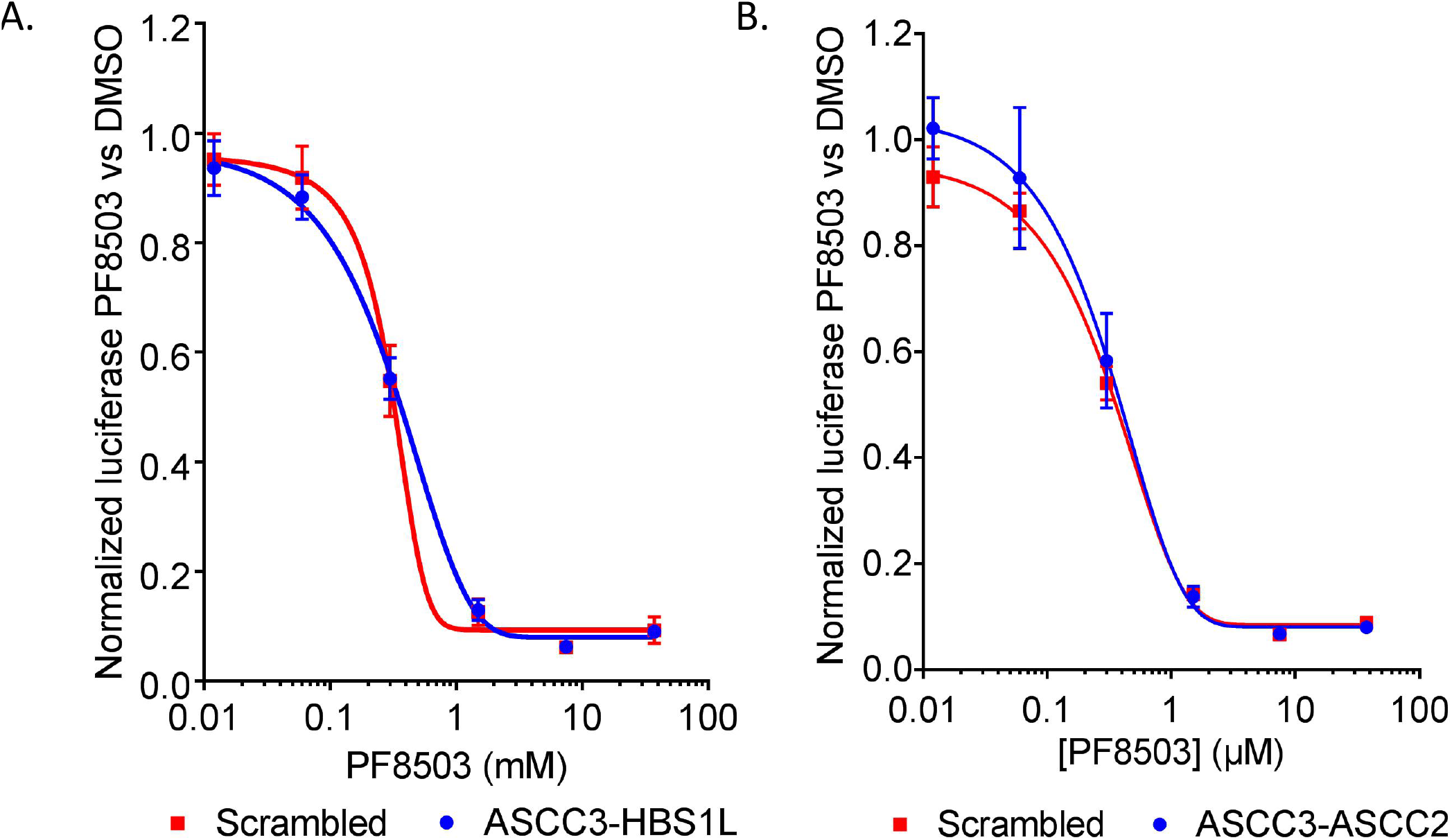
Effects of double knockdowns on PCSK9 reporter inhibition. Effect of PF8503 dose on the relative signal of PCSK9(1-35)-Rluc and control Fluc mRNA reporters after 7-8 hr incubation in K562 double-knockdown cell lines. These cell lines were obtained by using single lentiviral constructs as shown in Fig 5A, with mU6-HBS1L-hU6-ASCC3 (A) or mU6ASCC2-hU6ASCC3 (B). Average of triplicate experiments with standard deviation shown.

**S12 Fig.**
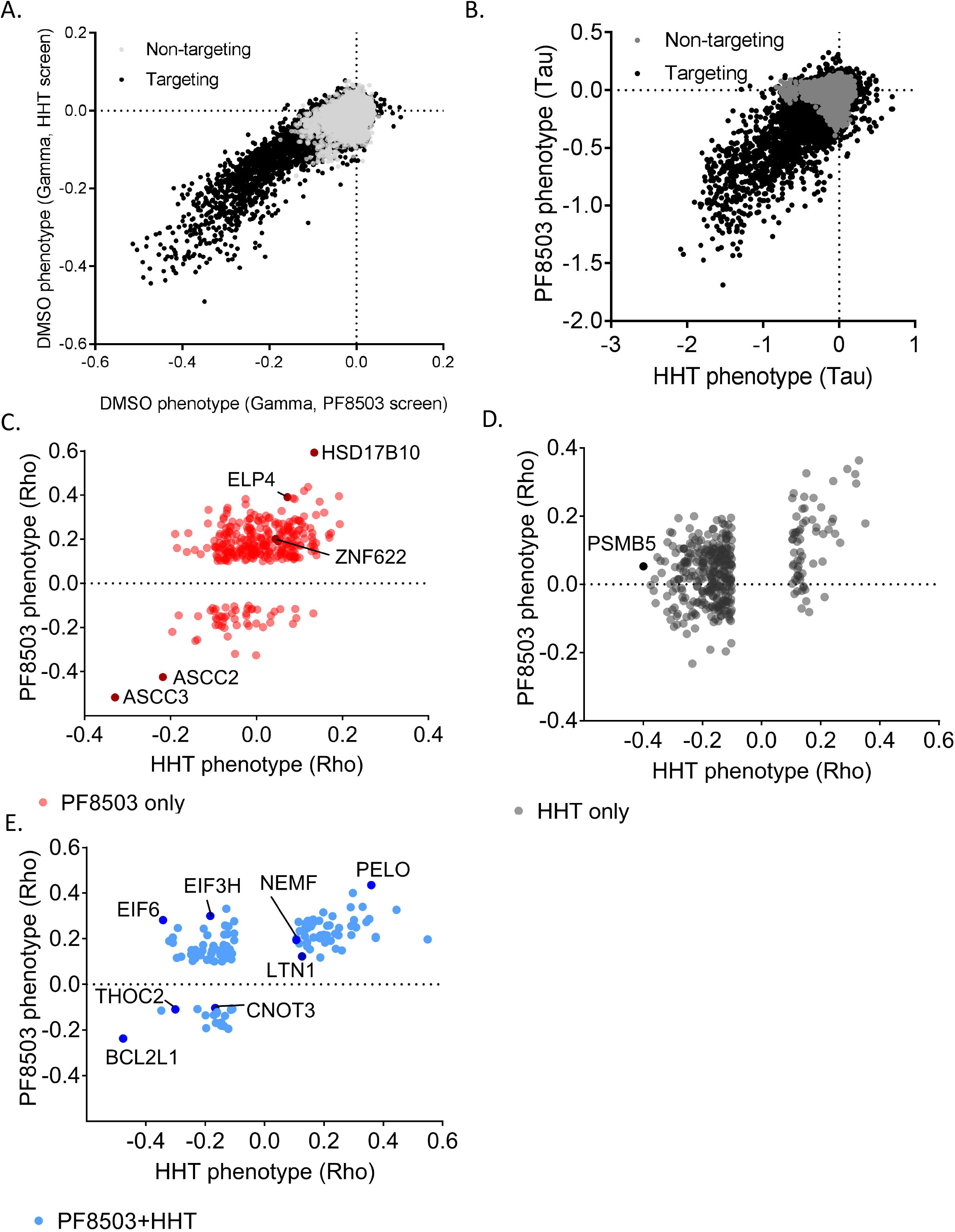
Comparisons between HHT and PF8503 CRISPRi screens. (A) Correlation between the untreated phenotypes (Gamma) obtained in the PF8503 and HHT screens, which were performed independently. (B) Correlation between the compound-treated phenotypes (Tau) obtained in the PF8503 and HHT screens. (C-E) Comparison of Rho phenotypes obtained in the HHT screen and in the PF8503 screen, for genes impacting cell fitness with PF8503 only (C), PF8503 and HHT (D), or HHT only (E).

**S1 Table. Proteins affected by PF8503 and PF846 as revealed by ribosome profiling.** *undef, undefined as no discrete stall site could be identified 5’ of the DMax position.

**S2 Table. Comparison of proteins affected by PF846 in Huh-7 cells in the present and previous ribosome profiling experiments.** *undef, undefined as no discrete stall site could be identified 5’ of the DMax position.

**S3 Table. CRISPRi results with PF8503.** *Transcript definition: P1, sgRNA targets transcription start site from P1 promoter as defined in the FANTOM5 data [46]; P2, sgRNA targets transcription start site from P2 promoter; P1P2, sgRNA targets transcription start site from P1 and P2.

**S4 Table. CRISPRi results with HHT**

**S5 Table. sgRNAs used in CRISPRi validation experiments**

**S6 Table. Antibodies used in Western blot analysis**

**S7 Table. Primers used in RT-qPCR experiments**

**S1 Text. Scripts used for Riboseq data analysis**

Additional file S1: Shell script used to perform alignment and filtering of raw sequences (without rRNA) and to create the nucleotide and codon maps.

Additional file S2: Python script used to filter alignments. Modified from [10]

Additional file S3: Positional_read_map.py, script used to make the nucleotide readmaps. Modified from [10]

Additional file S4: This script converts a nucleotide map to a codon map. Identical to

Additional file S5: This script normalizes a nucleotide or codon-based readmap to reads per million. Identical to [10]

Additional file S6: This script makes cumulative readmaps and DMax calculations. Identical to [10]

Additional file S7: This script filters CDS reads 3’ of the DMax position. Modified from [10].

